# Ahr-bacterial diet interaction modulates aging and associated pathologies in *C. elegans*

**DOI:** 10.1101/2020.04.13.021246

**Authors:** Vanessa Brinkmann, Alfonso Schiavi, Anjumara Shaik, Daniel R. Puchta, Natascia Ventura

## Abstract

Genetic and environmental factors concurrently shape the aging process. The nematode *Caenorhabditis elegans* has been instrumental in the past 30 years to identify most genes and interventions nowadays known to modulate aging (Lopez-Otin et al. 2013, Tigges et al. 2014, Dato et al. 2017). The aryl-hydrocarbon-receptor (AhR) was discovered as a dioxin-binding transcription factor involved in the metabolism of different environmental toxicants and xenobiotics (Poland et al. 1976). Since then, the variety of pathophysiological processes regulated by the AhR has rapidly grown, ranging from the immune response to cell death and metabolic pathways, and we disclosed it promotes aging phenotypes across species (Eckers et al. 2016, Esser et al. 2018). Interestingly, many AhR modulators may impact on aging and age-associated pathologies, but, whether their effects are actually AhR-dependent has never been explored. Here, we show for the first time that lack of *C. elegans* AHR-1 affects health and lifespan in a context-dependent manner. Using known mammalian AhR modulators we found that, similar to mammals, *ahr-1* has protective effects against environmental insults and identified a new role for AhR-bacterial diet interaction in animal lifespan, stress resistance, and age-associated pathologies. We narrowed down the dietary factor involved in the observed AhR-dependent features to a bacterially-extruded metabolite likely involved in tryptophan metabolism. This is the first study clearly establishing *C. elegans* as a good model organism to investigate evolutionarily conserved functions of AhR-modulators and -regulated processes, indicating it can be exploited to contribute to the discovery of novel information about AhR in mammals.

**Significance Statement:** The roundworm *C. elegans* has been instrumental for the identification of many of the genetic and environmental factors known to concurrently shape the aging process. The aryl-hydrocarbon-receptor (AhR) was originally discovered as a dioxin-binding transcription factor involved in the metabolism of environmental toxicants and, in the past, we found it promotes aging phenotypes across species. Many AhR modulators may impact on aging and age-associated pathologies, but, whether their effects are actually AhR-dependent has never been explored. We show here for the first time that *C. elegans* AhR has protective effects against environmental insults such as UVB radiations and the xenobiotic BaP and identified a new critical role for AhR-bacterial diet interaction in animal lifespan, stress resistance, and age-associated pathologies.

## Introduction

Aging affects every human and is accompanied by increased morbidity (*e*.*g*., diabetes, cardiovascular diseases, neurodegenerative diseases, and cancer) and risk of death (1, 2). In the past decades, also thanks to the growing number of researchers exploiting simple but powerful model organisms such as the nematode *Caenorhabditis elegans* (*C. elegans*), numerous hallmarks of aging as well as genetic and environmental factors regulating it have been identified (2-4).

The aryl hydrocarbon receptor (AhR) is a highly conserved transcription factor of the bHLH PAS family originally discovered as a dioxin-binding protein (5) and involved in the metabolism of different environmental toxicants and xenobiotics. Since its discovery, the variety of pathophysiological processes regulated by the AhR has grown and range from cell death, to immune response and neuronal development (6). Contradictory studies also indicated a role for AhR in the aging process (reviewed in (7)) and more recently, pro-aging functions of AhR (8, 9) have been described in an evolutionarily conserved manner from *C. elegans* to mammals. In mammals, in basal conditions, AhR is bound by HSP90, AIP, and p23, which retain it in the cytoplasm, in a ligand-affine state. Ligand binding of an AhR agonist leads to the dissociation of the AhR binding complex and AhR nuclear translocation (10, 11). In the nucleus, AhR dimerizes with the AhR nuclear translocator (Arnt), and the AhR-Arnt heterodimer then binds to the xenobiotic responsive elements (XREs) on AhR target genes (11). Similar to its mammalian counterpart, the *C. elegans* AhR homolog, AHR-1, forms a heterodimer with the *C. elegans* Arnt homolog AHA-1 and binds to XREs (12). It is expressed in several types of neurons and plays a role in neurodevelopment and long-chain unsaturated fatty acid synthesis (13-17). However, unlike mammalian AhR it does not bind to TCDD or □-naphthoflavone (12), but no other modulators have been tested so far for their effect on *C. elegans* AHR-1 activity and downstream effects. Although AhR was originally discovered as a dioxin-binding protein (5) in more recent years many other compounds have been identified which modulate its activity and can be mainly divided into four categories: xenobiotics (*e*.*g*., 2,3,7,8-tetrachlorodibenzodioxin (dioxin) and benzo(a)pyrene (BaP)), dietary factors (*e*.*g*., kaempferol, and curcumin), endogenous modulators (*e*.*g*., 6-formylindolo[3,2-b]carbazole (FICZ) and kynurenine) and ligands generated by the microbiota metabolism (*e*.*g*., indole-3-acetate and tryptamine) (5, 18-24). Interestingly, many of these AhR modulators may impact on aging and age-associated pathologies (25-27), but whether these effects are AhR-dependent and whether AhR itself influences the progression of these diseases has been largely unexplored. Here, using *C. elegans* as a powerful model organism for aging and toxicology studies, we followed up on our previous finding indicating an anti-aging effect of *C. elegans ahr-1* depletion (8) and investigated the effect of possible AHR-1 modulators on aging and other age-related features, such as stress response and associated pathologies. Our findings indicate that *C. elegans* can be exploited to study evolutionarily conserved functions of AhR modulators and their regulated processes and therefore to contribute to the discovery of novel information about AhR in mammals. Most importantly, we identify aging and associated pathologies as novel life traits regulated by the AhR in a diet-dependent manner bringing further complexity to the landscape of AhR-microbiota regulated processes.

## Results

### *C. elegans* AHR-1 is differentially involved in stress response

Loss of AHR-1 extends *C. elegans’* health- and lifespan (8). Since lifespan extension often correlates with increased resistance to different types of stressors (28), we assessed AhR role in stress resistance by exposing wild-type and *ahr-1* mutants to either heat shock, metabolic stressors (i.e. high concentrations of glucose or the hypoxia mimetic iron chelator 2,2’Bipyridyl, BP) or UVB radiations. As expected, the development and fertility of wild-type animals were affected by all insults in a dose-dependent manner (**Figure 1**) (29, 30). While consistent with our previous data (8) loss of *ahr-1* increased heat-stress resistance (**Figure 1 A, B**), it did not confer resistance to any of the newly tested insults (**Figure 1 C - H**). Actually, UVB-induced developmental delay and embryonic lethality were significantly more affected in the *ahr-1(ju145)* mutants than in wild-type worms (**Figure 1 G, H**).

**Figure 1.**
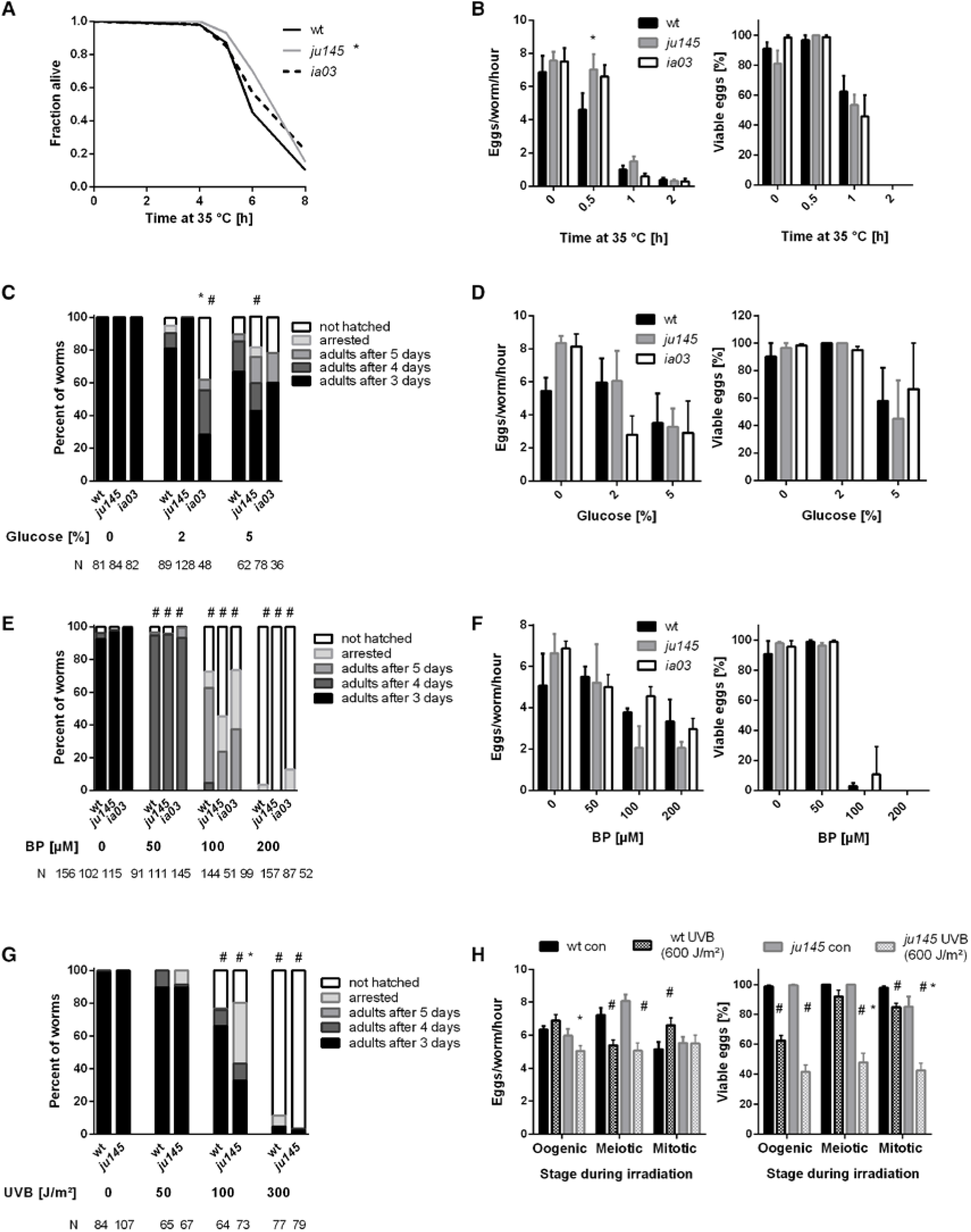
Loss of *ahr-1* differentially regulates resistance to stressors. **A)** Survival in response to heat shock. Curves show the pooled data of 60 worms in 3 independent replicates. Statistical test: Log-Rank test, * significance *vs*. wild-type. **B)** Fertility after heat stress. Shown are the number (left panel) and viability (right panel) of eggs laid from gravid adults treated with heat shock for the indicated time. Mean + SEM of pooled data from 18 worms/condition in 3 independent experiments are shown. **C - D)** Development and fertility in response to the indicated concentration of glucose. Means (+SEM) of 3 independent replicates are shown. N = number of individuals in panel C, 9 individuals were used in panel d. **E – F)** Development and fertility in response to the indicated concentration of iron chelator (BP). Means (+SEM) of 3 and 4 independent replicates are shown. N = number of individuals in panel E, 9 individuals were used in panel F. **G – H**) Development and fertility in response to indicated doses of UVB. Fertility was assessed at a dose of 600 J/m^2^. Means (+SEM) of 3 independent replicates are shown. N = number of individuals in panel G, 9 individuals were used in panel H. **B – H)** Statistical test: 2-way ANOVA with Tukey’s multiple comparisons test, * significance *vs*. wild-type, # significance *vs*. control (untreated), p-value < 0.05.

To further characterize animals’ stress response, we quantified the expression of *C. elegans* transgenic reporters for different phase-I and phase-II detoxification genes typically regulated by AhR in mammals: *cyps* (*e*.*g*., *CYP1A1* or *CYP1B1*), *ugts* (*e*.*g*., *UGT1A1* or *UGT1A6*) and *gsts* (*e*.*g*., *GSTA1* or *GSTA2*) (11, 31, 32). Out of nine tested reporters, we found five genes (*cyp-35A2, cyp-35B1, gst-4, cyp-37A1*, and *ugt-29*) differentially expressed by at least ten percent upon *ahr-1* RNAi (**Figure S1; Table S1**). Unexpectedly, when crossed into the *ahr-1(ju145)* mutant background, we observed different effects for *cyp-35B1* and *gst-4* expression, while the expression of *ugt-29* remained reduced in the *ahr-1* mutant (**Figure S1 C, D, E**), suggesting either different mode of action of *ahr-1* silencing and mutation or tissue-specific effects disclosed by the RNAi treatment. In line with the differential effect between *ahr-1* RNAi and the mutant, *ahr-1* RNAi did not affect the lifespan of the wild-type animals (**Table 1; Figure S1 G**). Most notably, while in most cases tissue-specific *ahr-1* RNAi had no effects, health- and lifespan were significantly shortened when RNAi was applied systemically and enhanced in the nervous system (**Table 1; Figure S1 F**). Taken together these data reveal that a complex AHR-1 role in *C. elegans’* lifespan and response to stress in a tissue- and insult-dependent manner.

**Table 1.**
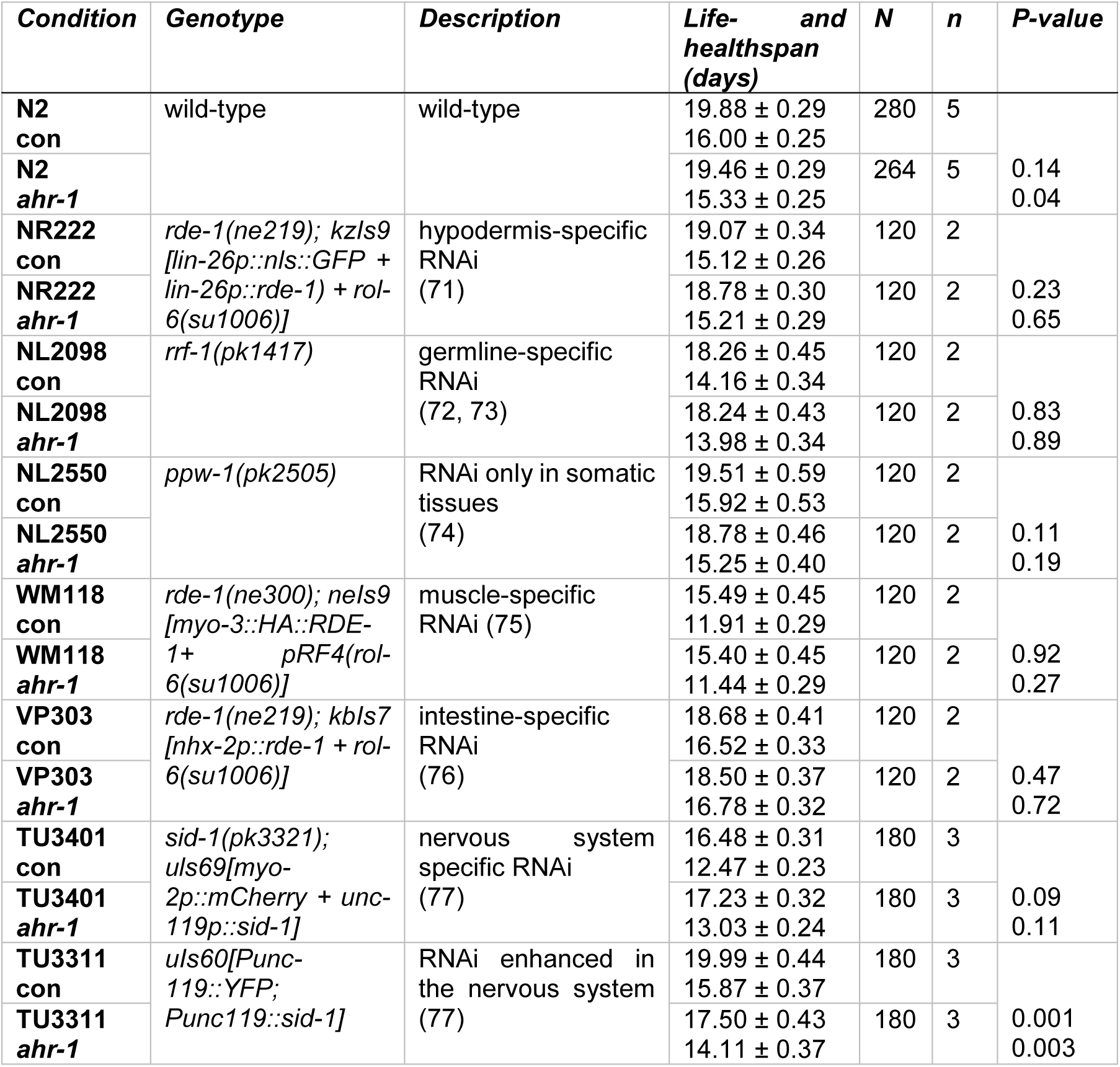
Survival analysis upon tissue-specific *ahr-1* RNAi. Mean life- and healthspans +/- SEM of pooled data are shown. The L4440 plasmid was used as a negative control. Statistical analysis was performed between *ahr-1* and control RNAi using Log-rank test. The number of worms is specified as N, while the number of replicates is specified as n.

### AHR-1 differentially regulates lifespan in response to potential modulators of its activity

In mammals, AhR exerts its functions primarily in response to ligands or modulators of its activity, such as exogenous substances (e.g. xenobiotics) or endogenous products of metabolism (e.g. microbiota-associated factors) (23). The xenobiotic BaP exerts toxic effects in mammals through AhR-dependent *cyps* expression (24), and it can induce *cyps* expression also in *C. elegans* (33). We observed that although BaP significantly increased the expression of *cyp-35B1* and affected animals’ development in a dose-dependent manner, these effects were largely *ahr-1*-independent (**Figure S2 A, B**). Interestingly, we found that BaP treatment from adulthood, consistent with its toxic effects, significantly shortened animals’ health- and lifespan, with a more pronounced effect on *ahr-1* mutants, indicating a protective role of AHR-1 against BaP-curtailed longevity (**Figure 2 A, B**). In mammals, 6-formylindolo[3,2-b]carbazole (FICZ), a photoproduct of tryptophan produced in response to UVB light, is another AhR high-affinity ligand (19, 20). In *C. elegans* UVB radiation shortens lifespan, induces embryonic lethality and germline cells apoptosis in a dose-dependent manner (34, 35). We showed that *ahr-1(ju145)* is more sensitive to UVB-induced embryonic lethality (**Figure 1 H**). Accordingly, we found that UVB-induced germline cells apoptosis is significantly more increased in the *ahr-1* mutant’s germline compared to wild-type animals (**Figure S2 C**). UVB irradiation increases *Cyp1A1* expression in human keratinocytes in an AHR-dependent manner (19). Interestingly, similar to the above results with BaP, although UVB tent to increase *cyp-35B1* expression in an *ahr-1*-independent manner (**Figure S2 D**), it decreased animals’ lifespan and motility with a significantly stronger effect on the *ahr-1(ju145)* compared to wild-type animals (**Figure 2 C, D**). These results agree with the notion that loss of AhR sensitizes mammalian cells to UVB-induced apoptosis (36) and support an evolutionarily conserved protective role of AhR in response to two of its most common activators.

**Figure 2.**
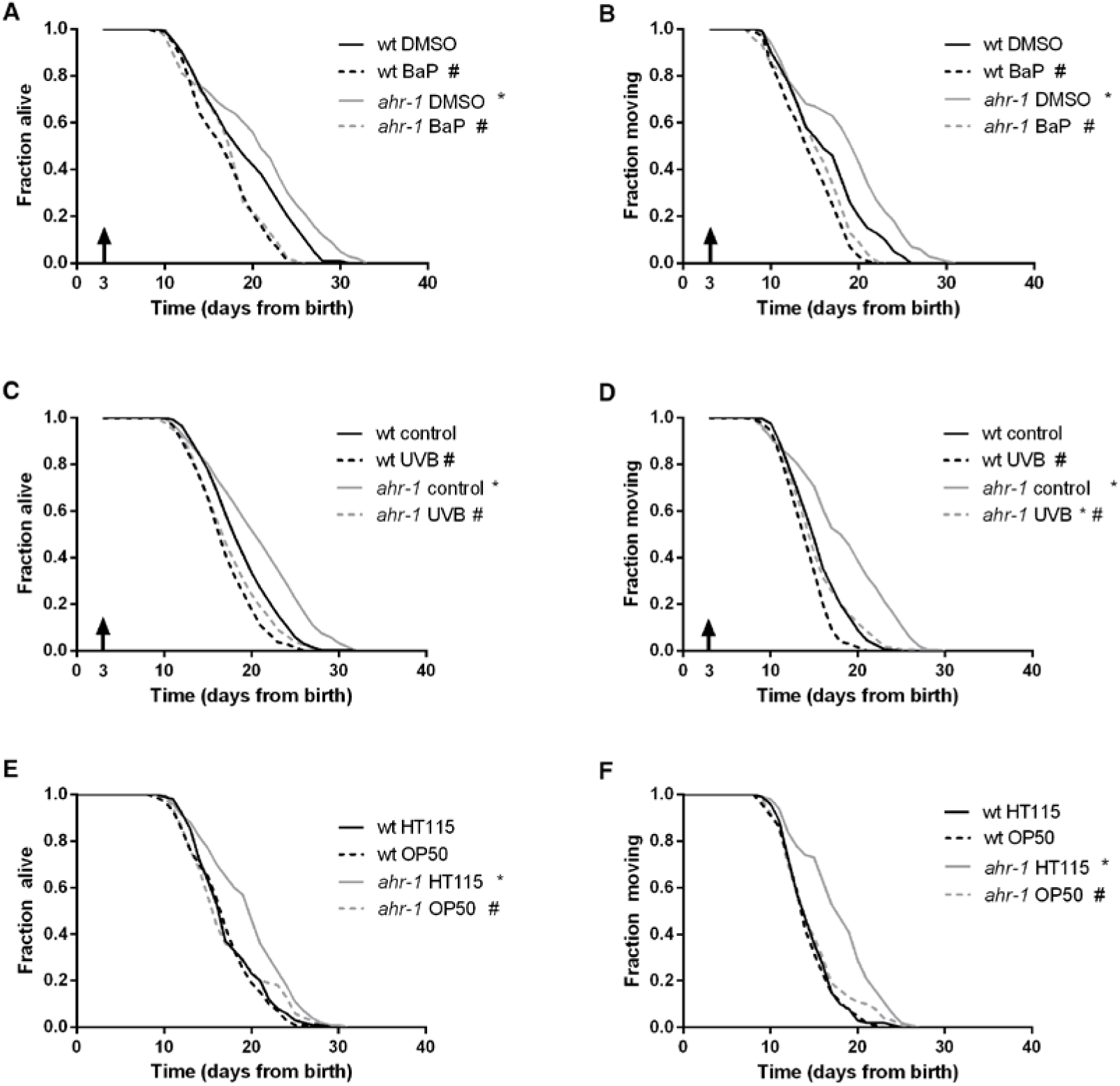
AHR-1 displays evolutionarily conserved functions and affects aging in a diet-dependent manner. **A – B)** *ahr-1* is more sensitive to xenobiotic stress. Pooled lifespan/healthspan curves of 120 worms/condition in 2 independent replicates treated with either DMSO or 5 µM BaP from adulthood are shown. **C – D)** *ahr-1* is more sensitive to UVB stress. Pooled lifespan/healthspan curves of 300 (wt control, *ahr-1* control), 180 (wt UVB) and 178 (*ahr-1* UVB) worms/condition in 3 independent replicates either left untreated or treated with 1200 J/m^2^ UVB from adulthood are shown. **E – F)** AHR-1 affects aging in a diet-dependent manner. Pooled lifespan/healthspan curves of 170 (wt OP50) and 180 (all other conditions) worms/condition in 3 independent replicates grown either on HT115 or OP50 are shown. **A – F)** Statistical test: Log-Rank test, # significance *vs*. control/HT115, * significance *vs*. wt, p-value < 0.05.

Finally, as microbiota-associated factors modulate AhR activity in mammals, we took advantage of two common *Escherichia coli* strains used as a food source for *C. elegans*, HT115(DE3) and OP50(xu363). Strikingly, we found that the reduced expression of the *cyp-35B1*::GFP reporter as well as the beneficial effects on lifespan, motility, pharyngeal pumping, and heat resistance elicited in the *ahr-1* mutants fed HT115 are abolished when animals are fed OP50 (**Figure 2 E, F; Figure S2 E - G**). These differences are neither due to extrinsic bacterial differences nor to the effect of the bacteria on the *ahr-1* expression (**Figure S2 H**) thus likely relying on the modulation of AHR-1-regulated processes. Results described so far indicate that known AhR modulators differentially affect *C. elegans* lifespan in an *ahr-1*-dependent manner but that classical detoxification related genes (*cyp*s) are possibly not the major targets of *C. elegans* AHR-1. Thus, to gain further insight into molecular mechanisms of AhR-regulated stress response and aging we turned to the results of a transcriptomic analysis recently carried out in the lab (Brinkmann et al. *in preparation*). A thorough analysis of the most differentially expressed genes between wild-type and *ahr-1(ju145)* revealed that many of them have been described in *C. elegans* to be affected not only during aging but also by dietary compounds (e.g. quercetin, resveratrol, bacteria) known to modulate AhR activity in mammals (**Tables S2, S3**). Very interestingly, quantitative PCR analysis of some of these genes (*atf-2, K04H4*.*2, egl-46, T20F5*.*4, ptr-4, dyf-7, clec-209, C01B4*.*6, C01B4*.*7, F56A4*.*3*) confirmed their *ahr-1* dependency in basal conditions, and identified a specific AhR-bacteria diet regulatory effect: their expression was generally not affected by UVB or BaP but their reduced expression in the *ahr-1(ju145)* mutants fed H115 bacteria, similar to *cyp-35B1*, was abolished in animal fed and OP50 diet (**Figure S3)**. Altogether, results shown so far support an evolutionarily conserved protective role for AhR against BaP and UVB, and identify a new role for AhR in environmentally regulated aging with dietary bacteria as an important component in *ahr-1-*signaling-mediated longevity.

### Loss of *ahr-1* affects age-associated pathologies in a diet-dependent manner

The critical role of AHR-1 in *C. elegans* neurons (14, 15, 17) and motility (8, 9) prompted us to investigate whether diet-dependent effects also influence age-related neuromuscular pathologies. A growing body of evidence indeed indicates microbiota can affect various aspects of *C. elegans’* health-span (37-41). Of note, in line with the results on the wild-type strain, loss of *ahr-1* extended life- and health-span of two age-associated disease models, namely animals with muscle expression of aggregation-prone proteins polyQ_40_ (42) and α-synuclein (43) (**Figure 3**), and these beneficial effects were significantly suppressed when animals were fed an OP50 diet instead of HT115 (**Figure 3 A, B, E, F**). Despite being longer-lived, *ahr-1* mutants fed HT115 displayed a greater number and size of polyQ_40_ and α-synuclein aggregates, which is nonetheless in line with studies revealing no direct correlation between aggregates and toxic effects (44). Moreover, the increase in aggregation was mainly regulated in a diet-independent manner (**Figure 3 C, D, G, H**), indicating that the *ahr-1*-diet interaction effect on health-span is independent of its effect on protein aggregation. Somewhat surprisingly, when we investigated the effect of *ahr-1* deficiency in another pro-aggregation model - a *C. elegans* strain expressing a pan-neuronal human Aβ peptide (45) - we found that contrary to the other models, it reduced animals’ life- and health-span, yet interestingly in a diet-dependent manner (**Figure S4**).

**Figure 3.**
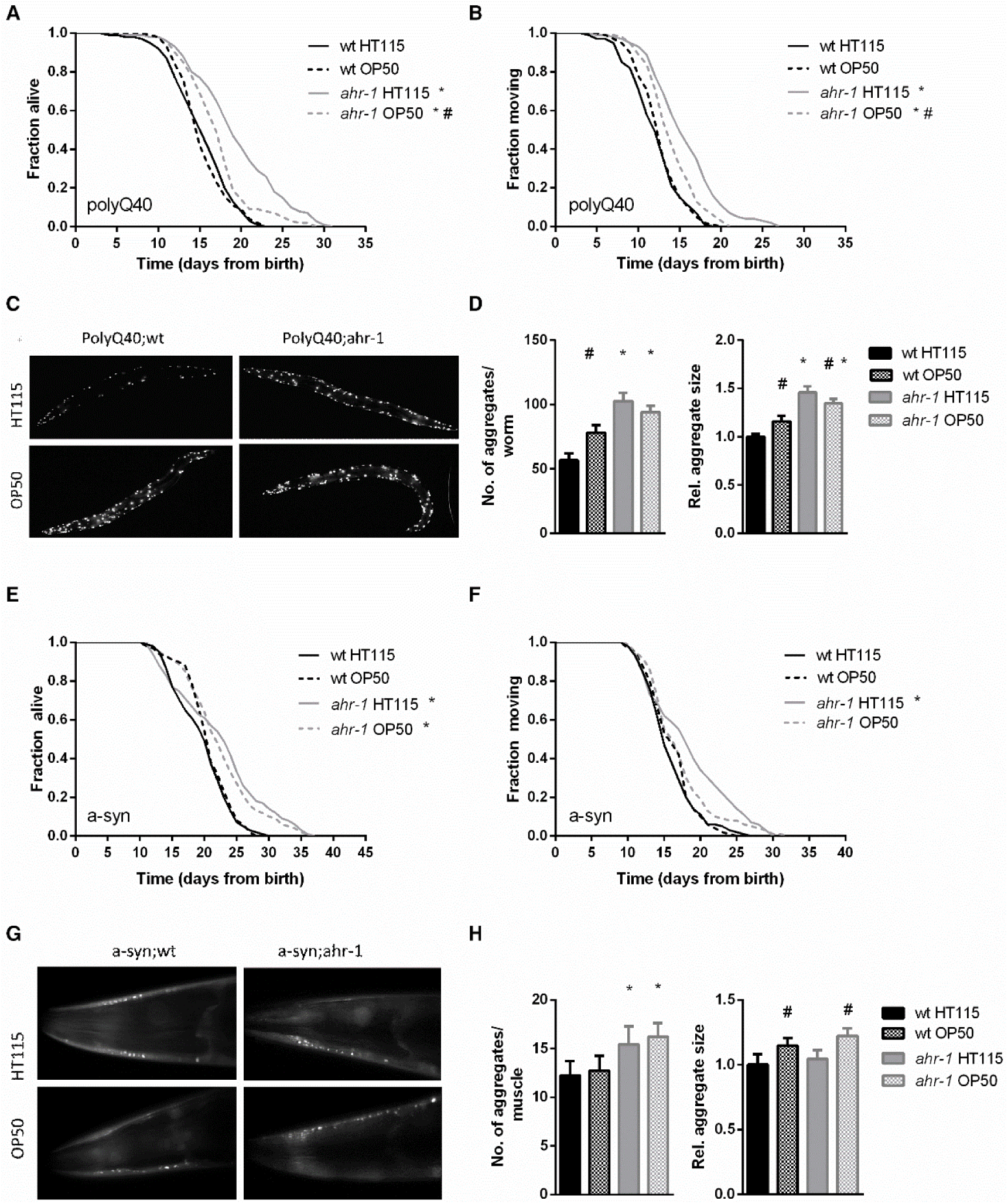
AHR-1 mutants increase aggregation but extend lifespan in a diet-dependent manner. **A – B)** Kaplan Meier curves of polyQ;wt and polyQ;ahr-1 of 180 worms/condition in 3 independent experiments are shown. * p-value < 0.05 *vs*. polyQ;wt, # p-value < 0.05 *vs*. HT115, statistical test: Log-rank test. **C)** Representative fluorescence images of 10-days old polyQ;wt and polyQ;ahr-1 on HT115 and OP50. **D)** Quantification of aggregates in 10-days old polyQ;wt and polyQ;ahr-1. Mean + 95 % CI of pooled data from 34 (wt HT115), 29 (wt OP50), 26 (*ahr-1* HT115), and 35 (*ahr-1* OP50) worms in 3 independent replicates is shown. Statistical test: One-way ANOVA with Tukey’s multiple comparisons test, * p-value < 0.05 *vs*. polyQ;wt, ^#^ p-value < 0.05 *vs*. HT115. **E – F)** Kaplan Meier curves of a-syn;wt and a-syn;ahr-1 of 120 worms/condition in 2 independent experiments are shown. * p-value < 0.05 *vs*. a-syn;wt, # p-value < 0.05 *vs*. HT115, statistical test: Log-rank test. **G)** Representative fluorescence images of the head muscles of 7-days old a-syn;wt and a-syn;ahr-1 on HT115 and OP50. **H)** Quantification of aggregates in 7-days old a-syn;wt and a-syn;ahr-1. Mean + 95 % CI of pooled data from 77 (wt HT115), 82 (wt OP50), 88 (*ahr-1* HT115), and 92 (*ahr-1* OP50) worms in 3 independent replicates is shown. Statistical test: One-way ANOVA with Tukey’s multiple comparisons test, * p-value < 0.05 *vs*. a-syn;wt, ^#^ p-value < 0.05 *vs*. HT115.

Given that temperature may affect aging and associated pathologies depending on the bacterial diet and genetic background (37, 46) we next tested whether the healthy aging phenotype of *ahr-1* is also temperature-dependent. Despite the increased resistance to heat shock of *ahr-1(ju145)* mutants, and opposite to the lifespan at 20 °C, *ahr-1*-depleted animals were short-lived on an HT115 diet at 25 °C. Yet, again, this difference in the lifespan of wild-type and *ahr-1* was abolished on the OP50 diet, which *per se* already shortened the lifespan at 25 °C of wild-type animals (**Figure 4 A, B**). The higher temperature also prevented the beneficial diet-dependent effects of loss of *ahr-1* on life- and health-span in the polyQ strain (**Figure 4 C, D**), while the number of polyQ aggregates increased according to the rise in temperature but still in a diet-independent manner (**Figure 4 E, F**). Overall our data uncovered a new diet-dependent effect of AhR in modulating aging and associated neuromuscular pathologies, which is nonetheless uncoupled from age-dependent increase in protein aggregation.

**Figure 4.**
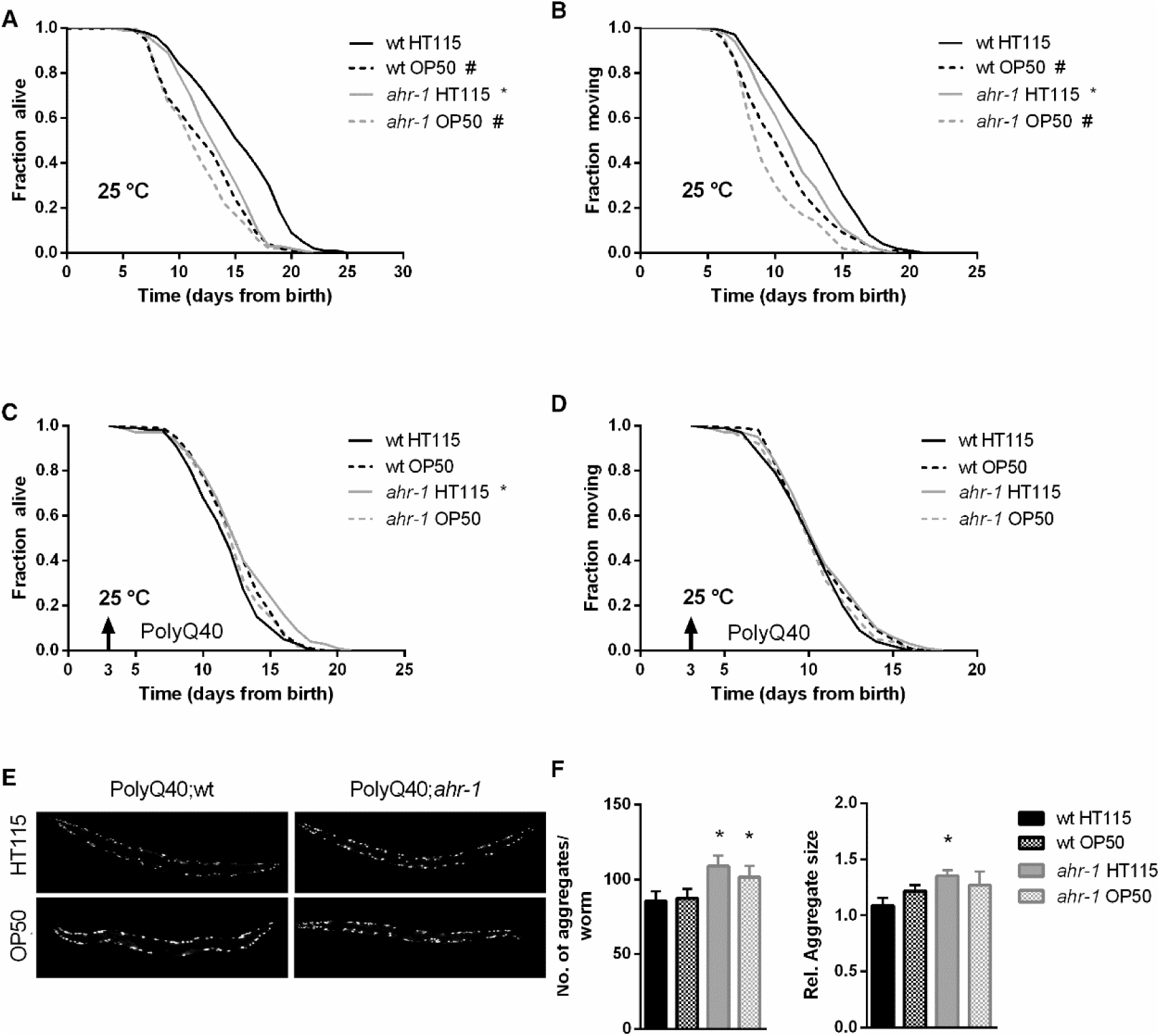
At 25 °C, *ahr-1(ju145)* is short-lived and loses its protection against the toxicity of polyQ_40_ aggregates. **A – B)** Kaplan Meier curves of wild-type and *ahr-1(ju145)* at 25 °C. Pooled data of 180 worms/condition in 3 independent replicates are shown. * p-value < 0.05 *vs*. wt, # p-value < 0.05 *vs*. HT115, statistical test: Log-rank test. **C – D)** Kaplan Meier curves of polyQ;wt and polyQ;ahr-1. Worms were grown at 25 °C from day 3 (indicated by arrowhead). Pooled data of 150 worms/condition in 3 independent replicates are shown. * p-value < 0.05 *vs*. wt, # p-value < 0.05 *vs*. HT115, statistical test: Log-rank test. **E)** Representative fluorescence images of 10-days old polyQ;wt and polyQ;ahr-1 grown at 25 °C from day 3. **F)** Quantification of aggregates in 10-days old polyQ;wt and polyQ;ahr-1 grown at 25 °C from day 3. Mean + 95 % CI of pooled data from 21 (wt HT115), 19 (wt OP50), 20 (*ahr-1* HT115) and 19 (*ahr-1* OP50) worms/condition in 2 independent replicates are shown. * p-value < 0.05 *vs*. wt, statistical test: One-way ANOVA with Tukey’s multiple comparisons test.

### Bacterial tryptophan metabolism mediates the beneficial effect of the *ahr-1* mutants

In search of the potential bacterial factor responsible for the diet-dependent effects, we first asked whether metabolically active bacteria are required for the observed differences. To this end, we compared the heat-stress resistance of animals fed alive bacteria with that of animals fed bacteria either killed before seeding on plates or killed on the plates two days after seeding (thus allowing metabolites secretion). Very interestingly, killing bacteria before seeding completely abolished the differences between wild-type and *ahr-1* fed HT115 (**Figure 5 A - B**), indicating that a factor produced by metabolically active HT115 bacteria may influence the AHR-1-mediated effects. In support of this possibility, the increased resistance to stress of *ahr-1* mutants observed on living HT115 bacteria persisted when bacteria were killed after growing for two days on the feeding plates (**Figure 5 C**). In line with the heat shock experiments, killing the bacteria before seeding also completely suppressed the increased lifespan of the *ahr-1* mutants on HT115 with no major effects on animals fed OP50 (**Figure 5 D**). These data point towards HT115 secreted metabolites playing a role in *ahr-1*-mediated life- and health-span, likely via gut ingestion or neuronal sensing (37). A mass spectrometric analysis of the two bacteria supernatants revealed that the subtracted spectrum between HT115 and OP50 showed three prominent peaks (**Figure 5 E**) with m/z 361 likely being an arginine-tryptophan dipeptide. On the other hand, the medium of OP50 was enriched in an alanine-glutamate dipeptide (**Figure S5**). In mammals, tryptophan (Trp) and its metabolites (such as indole) modulate AhR activity (22, 47, 48) and in *C. elegans* Trp was shown to abolish some of the different phenotypes observed between alive HT115 and OP50 (49). Of note, supplementation of L-tryptophan completely abolished the heat-stress resistance of *ahr-1(ju145)* on HT115 - and partially suppressed the difference also between wild-type and *ahr-1(ju145)* on OP50 (**Figure 5 F, G**). Overall, these data point towards a potential role in Trp metabolism in *ahr-1-*bacteria-regulated age-associated phenotypes.

**Figure 5.**
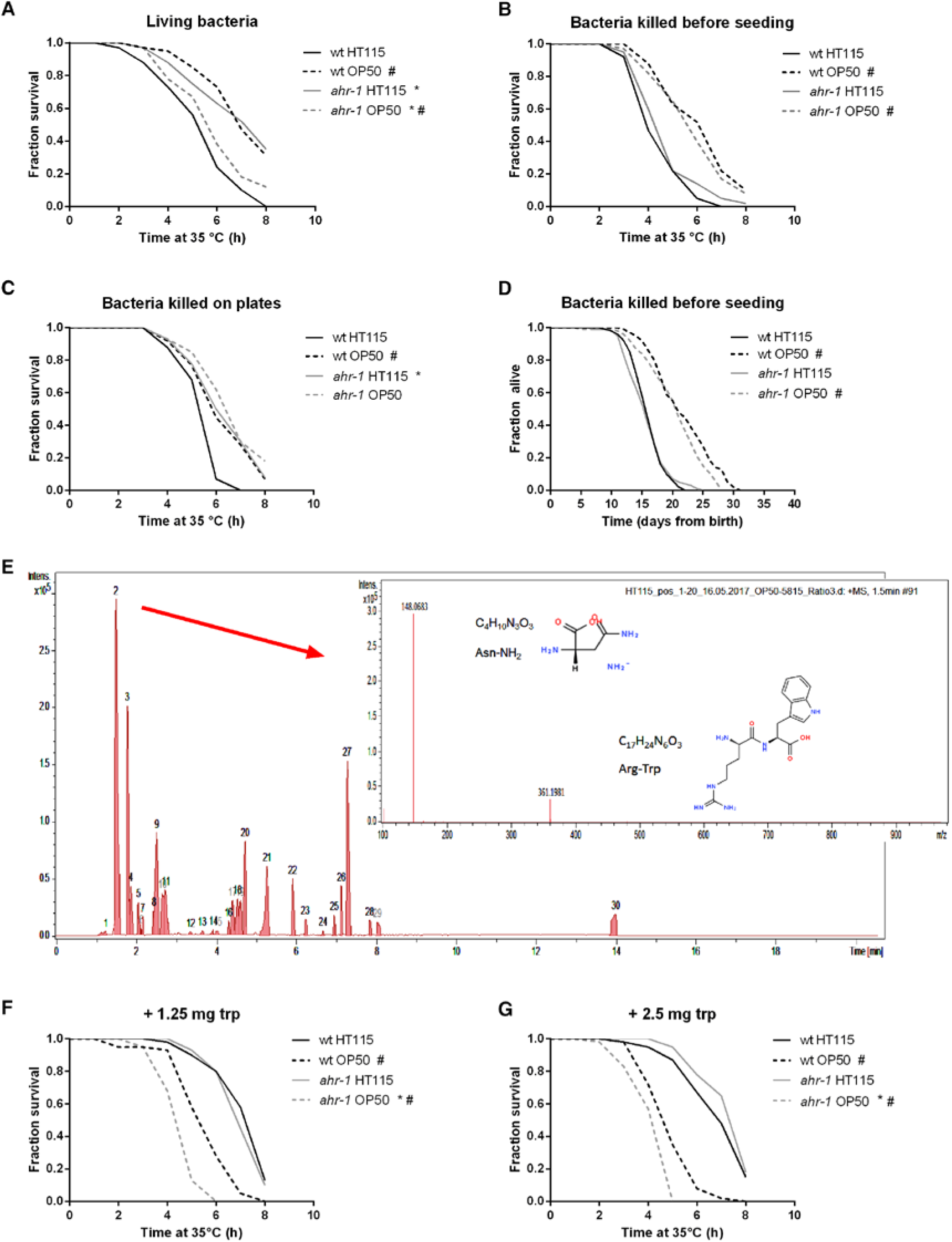
Metabolically active bacteria are required for the differences in lifespan between wild-type and *ahr-1* on HT115. **A – C)** Survival upon heat stress in 7-days old wild-type and *ahr-1(ju145)* feeding on either HT115 or OP50. Pooled data of 60 worms/condition in 3 independent experiments are shown. * p-value < 0.05 *vs*. wt, # p-value < 0.05 *vs*. HT115, statistical test: Log-rank test. **A)** Living bacteria were used as a food source. **B)** Bacteria were killed by UVB irradiation before seeding to the NGM. **C)** Bacteria had grown on the NGM for 2 days before being killed by UVB irradiation. **D)** Kaplan Meier curves of wild-type and *ahr-1(ju145)* on UVB-killed bacteria. Pooled data of 120 (wt HT115, *ahr-1* HT115, *ahr-1* OP50) and 110 (wt OP50) worms in 2 independent experiments are shown. * p-value < 0.05 *vs*. wt, # p-value < 0.05 *vs*. HT115, statistical test: Log-rank test. **E)** Positive ESI MS analysis. The HT115 BPC after the subtraction of the OP50 BPC is shown. Masses in peak 2 are shown as inset. **F – G)** Heat stress survival after tryptophan supplementation with indicated concentrations of tryptophan. Survival curves of 7-days old worms feeding on HT115 supplemented with tryptophan are shown. The curves show pooled data from or 40 worms/condition in 2 independent replicates (1.25 mg trp) or 60 worms/condition in 3 independent experiments (2.5 mg trp). * p-value < 0.05 *vs*. wt, # p-value < 0.05 *vs*. HT115, statistical test: Log-rank test.

## Discussion

*C. elegans* AHR-1, similar to mammalian AhR, forms a heterodimer with the *C. elegans* Arnt homolog AHA-1 and binds to xenobiotic responsive elements (XREs), but unlike its mammalian counterpart, it does not bind the classical AhR activators such as TCDD or □-naphthoflavone (12). However, no other potential AhR ligands or ligand-independent modulators have ever been tested in *C. elegans*, thus precluding to fully exploit the versatility of this model organism to unravel conserved or novel biological function for this environmentally relevant transcription factor. Only recently, conserved functions of the *C. elegans* AHR-1 in health and lifespan (8, 9) have been identified.

In this new study, we revealed a more general and complex role for this transcription factor in stress response and aging in basal conditions and in response to known modulators of its activity (i.e. curcumin, UVB and BaP). The differential effects on lifespan and gene expression we observed upon *ahr-1* genetic- or RNAi-mediated suppression interestingly reveal the importance of fine-tuning its activity in a dose and/or tissue-dependent manner, which very nicely recapitulate the variety of phenotypic features described in mammals upon dosage effects or tissue-specific depletion (7). Moreover, to our surprise, we found that besides increased resistance to heat-shock, lack of *ahr-1* did not confer resistance to any other investigated insult, such as metabolic stress or radiation, and in fact sensitize animals to UVB. We specifically decided to focus our attention on the influence on stress response and aging of different classes of mammalian AhR modulators: environmental (xenobiotic), endogenous (UVB induced FICZ) and dietary (microbiota) factors. These were shown to mainly induce AhR’s transcriptional activity in a ligand-dependent manner, but ligand-independent, as well as antagonistic functions, have been also suggested (11). Our data showed that *C. elegans ahr-1(ju145)* mutants are more sensitive to the lifespan shortening effects of UVB and BaP, two classical activators of mammalian AhR, thus further supporting a role for *ahr-1* in the aging process. Of note, *ahr-1* mutants also conferred sensitivity to UVB-reduced fertility, which we hypothesized to reflect increased sensitivity to germ cells apoptosis in irradiated animals as also found in human keratinocytes and mice (36, 47). Indeed, loss of AHR-1 function increased apoptosis in basal condition and upon radiation indicating a conserved anti-apoptotic function of the AhR in response to UVB. Similar to the detrimental effect of UVB, exposure to BaP in mammals causes a variety of cancers as well as neurotoxicity (48, 49), and loss of AhR prevents BaP- and UVB-induced carcinogenicity in mice (24, 47). In this study, we showed for the first time that BaP has a conserved detrimental effect on lifespan, which is significantly worsened in the *C. elegans ahr-1* mutants. Along with the UVB data, these findings support a protective role of AHR-1 in response to classical mammalian activators. However, surprisingly, both stressors induced *cyp-35B1* expression in an *ahr-1*-independent manner thus pointing to a non-canonical mode of activation of AHR-1 in *C. elegans*. Interestingly, BaP and the UVB-generated ligand FICZ are big and planar molecules which, based on preliminary *in silico* analysis (Brinkmann et al. *in preparation*) likely do not fit into the ligand binding pocket of the *C. elegans* AHR-1. Thus, one could speculate that AHR-1, like its mammalian homolog, is activated by reactive oxygen species (ROS) (50), which can be produced either by UVB (51) or BaP (52, 53).

Most notably, we identified for the first time a critical role for AHR-1-bacterial diet interaction in regulating aging and associated phenotypes. Specifically, *ahr-1* mutants displayed an extended health- and life-span on HT115 but not on the OP50 bacterial diet. Loss of *ahr-1* also ameliorated health- and life-span in models of age-associated pathologies with aggregation-prone proteins expressed in the muscles in a diet-dependent manner. Yet, surprisingly, the beneficial effect did not correlate with the amount and size of the aggregates, indicating either that aggregation in this context actually has a beneficial effect, or that loss of *ahr-1* promotes health-span independently from mechanisms regulating proteotoxic aggregation. Regardless, we hypothesized that bacteria-associated factors from HT115 or OP50 might mediate the diet-dependent changes in health-span and narrowed them down to secreted bacterial metabolites. Although the exact metabolite(s) responsible for the diet-dependent effects still remain to be identified, our data point to the potential involvement of Trp metabolism. This again supports evolutionarily conserved functions of AHR-1, as its activity is known to be influenced by Trp and its metabolites in mammals (22, 54, 55). Interestingly, Trp and its metabolites have been also shown to modulate *C. elegans’* health- and life-span. Trp supplementation increases heat-stress resistance and lifespan of *C. elegans* (56) and reduces the proteotoxicity in neurodegenerative disease models (44). Instead, different Trp metabolites display opposite effects: Trp degradation through the Kynurenine pathway increases proteotoxicity in neurodegenerative disease models (44), while the tryptophan metabolite indole, from commensal bacteria, increases the lifespan of *C. elegans* through AHR-1 (57).

Our analysis on bacterial supernatant also revealed differences in alanine-glutamate levels. Interestingly, it was recently shown that HT115, differently from OP50, possess the glutamate decarboxylate enzyme necessary to convert glutamate into GABA, which is ultimately responsible to protect *C. elegans* neurons from degeneration (59).Moreover, compared to HT115 fed worms, OP50 confer sensitivity to oxidative stress possibly due to mitochondrial alteration and increased ROS production (38, 59, 60). It will be interesting to assess whether glutamate and/or ROS metabolism play a role in bacteria-AHR-1 regulation of the aging process. The primary site of action of the bacterial metabolite (e.g. intestine, sensory neurons) as well as the underlying AhR-dependent molecular process (e.g. immune response, redox reactions), are also attractive aspects which remain to be elucidated

In summary, we demonstrated that *C. elegans ahr-1* displays evolutionarily conserved functions, such as its protective activity against BaP and UVB, and identified a new direct link between *ahr-1*-regulated processes and bacterial diet as a key determinant of aging and associated pathologies, most likely through tryptophan metabolism. Overall our findings support a central role for AhR in the aging process in a context-dependent manner, thus expanding the already vast panel of activities played by the AhR in different pathophysiological conditions and establish for the first time *C. elegans* as a powerful model organism to unravel new AhR-regulated processes in response to conserved modulators of its activity.

## Materials and Methods

### *C. elegans* strains and cultivation

*C. elegans* strains used in this study are listed in Table S4. We created the following strains for this study by crossing CZ2485 with different transgenic strains to obtain: NV33a: *ahr-1(ju145); cyp-35B1p::GFP + gcy-7p::GFP*, NV35a: *ahr-1(ju145); (pAF15)gst-4p::GFP::NLS*, NV38b: *ahr-1(ju145); unc-54p::Q40::YFP*, NV42a: *ahr-1(ju145); unc-54p::alpha-synuclein::YFP*,, NV47a: *ahr-1(ju145*); *ugt-29*p::GFP. For maintenance, worms were kept synchronized by egg lay at 20 °C on Nematode Growth Media (NGM) plates and fed with *E. coli* OP50 according to methods described in (61). For the experiments, worms were synchronized on plates supplemented with *E. coli* HT115(L4440) or OP50(L4440) according to the condition of interest. NGM plates were supplemented with 1 mM IPTG when using HT115(L4440) or OP50(L4440).

### Gene silencing by RNA-mediated interference (RNAi)

Gene silencing was achieved through feeding *E. coli* HT115(DE3) expressing plasmids with dsRNA against specific genes (62). RNAi feeding was applied continuously from birth to death.

### *E. coli* strains and growth

Bacteria were grown in LB medium at 37 °C overnight. When using *E. coli* carrying vectors the LB medium was supplemented with 0.01 % of ampicillin and 0.0005 % of tetracycline. *E. coli* HT115(L4440), HT115(*ahr-1*), and OP50 were obtained from the Ahringer *C. elegans* RNAi library (63). *E. coli* OP50*(xu363)* (40) was a gift from Shawn Xu.

### Heat stress survival

The resistance to heat stress was tested with 20 animals/condition per experiment at 35 °C on 3 cm plates wrapped with parafilm in an incubator (Intrafors HT Multitron). The number of dead animals was scored hourly by gently touching the worms with a platinum wire and analysis was performed as described for the lifespan assay.

### Reproduction on heat stress, glucose, and BP

Animals were grown on control plates with alive bacteria until day 3 and then transferred to treatment or control plates for 24 hours. After a 24-hour treatment, 3 animals of each condition were transferred to fresh control plates for 4 hours to lay eggs and the number of eggs laid was counted. The number of progenies hatched from these eggs was counted 2 days afterward.

### Development

The development on the specific compound was explored by counting the number of eggs, which developed to gravid adults after 72, 96, and 120 hours as well as the number of worms that arrested their development and the number of eggs, which did not hatch.

### Glucose treatment

D-Glucose (Merck, 8342) was dissolved in ddH_2_O and supplemented to the NGM after autoclaving and UVB killed *E. coli* HT115(L4440) were fed as a food source.

### 2,2′-dipyridyl (BP) treatment

The iron chelator 2,2′-dipyridyl (Carl Roth, 4153) was dissolved in ddH_2_O and supplemented to the NGM after autoclaving.

### UVB irradiation

Worms were exposed to ultraviolet radiation on bacteria-free NGM plates, using a Waldmann UV 236 B (UV6) lamp with an emission maximum of 320 nm. Irradiation times were 9 seconds, 27 seconds, and 53 seconds for 100 J/m^2^, 300 J/m^2^ and 600 J/m^2^, respectively with a distance of 18 cm between lamp and plate.

### Reproduction after UVB treatment

3 days old worms were treated with UVB and the effect of irradiation on germ-cells in different stages (oogenic stage, meiotic stage, and mitotic stage) was investigated, as described in (64). Briefly, the number and viability of eggs laid between 1 – 8 hours (oogenic stage), 8 – 24 hours (meiotic stage) and 24 – 32 hours (mitotic stage) after the irradiation were analyzed.

### Lifespan

The lifespan analysis was started from a synchronized population of worms, which was transferred to fresh NGM plates daily during the fertile period. After the fertile phase, the animals were transferred every alternate day. Dead, alive and censored animals were scored. Animals with internal hatching (bags), an exploded vulva or which died desiccated on the wall were censored. Survival analysis was performed in OASIS (65) or OASIS 2 (66) using the Kaplan Meier estimator. A log-rank test between pooled populations of animals was used for the evaluation of statistical significance. The p-values were corrected for multiple comparisons using the Bonferroni method.

### Movement/Healthspan

The movement was set as a parameter for healthy aging, and the phase of active movement is referred to as healthspan. It was assessed in the populations used for the lifespan assay. Animals, which were either crawling spontaneously or after a manual stimulus, were considered as moving while dead animals or animals without crawling behavior were considered as not moving. Statistical analysis was done as described for lifespan.

### Benzo(a)pyrene (BaP) treatment

Benzo(a)pyrene (Sigma Aldrich, B1760) was dissolved in DMSO (Carl Roth, 4720) in concentrations 1000 times higher than the desired concentration in the NGM. After autoclaving the NGM, BaP or DMSO were added to the media. For development assays, worms were treated from eggs, while they were treated from the first day of adulthood for lifespan and healthspan assays.

### Quantification of polyQ aggregates

PolyQ_40_ aggregates were visualized by fluorescence microscopy (100x magnification) in worms anesthetized with 15 mM sodium azide (Sigma, S2002). The number and the size of the aggregates were quantified in Fiji (67). To assess the number of aggregates, images were stitched using the Fiji pairwise stitching plugin (68) to create whole worms. The average size of the aggregates was instead measured in the non-stitched images. The number and the size of aggregates were counted using the plugin “Analyze Particles”.

### Quantification of α-synuclein aggregates

α-synuclein aggregates in the head muscles of 7-days old worms were visualized by fluorescence microscopy (400x magnification) in worms anesthetized with 15 mM sodium azide (Sigma, S2002). Pictures were segmented using Ilastik (version 1.3.0) (available on https://www.ilastik.org/) (69). The segmented pictures were used to analyze the number and size of the aggregates in Fiji (67) using the plugin “Analyze Particles”.

### Assessment of age-associated features at 25 °C

Lifespan, movement, and polyQ aggregation analysis were performed as described above. PolyQ-expressing worms were kept at 20 °C until reaching the L4 stage and were afterward kept at 25 °C for the rest of their lifespan.

### Killing bacteria before seeding

When seeding killed *E. coli* onto the NGM, the bacteria were pelleted (10 min at 4000 rpm), the supernatant was removed, and the pellet was suspended in S-basal to a final concentration of OD_595_ = 3.6. Then the bacteria suspension was irradiated with a UVB lamp (Waldmann UV 236 B) for 1 hour to kill the bacteria. The suspension of the dead bacteria was again pelleted and re-suspended in fresh S-basal. Killed bacteria were seeded to NGM plates in a concentration of OD_595_ = 3.6 and let dry at room temperature overnight.

### Killing bacteria on plates

Bacteria (OD_595_ = 0.9) were seeded to the NGM plates and let grow for 2 days at room temperature. The bacterial lawn was then killed by exposure to UVB light (Waldmann UV 236 B) for 45 min.

### Tryptophan supplementation

The concentration of tryptophan and the supplementation procedure was taken from (70) with minor changes. L-tryptophan (Carl Roth 4858.2) was dissolved in water at a concentration of 12.5 mg/ml and incubated shaking at 30°C for 45 min. Afterward, the solution was filtered (pore size: 0.22 µm, Carl Roth P666.1). For 7 ml of NGM 200 µl of 12.5 mg/ml tryptophan was spotted on the bacterial lawn of an NGM plate. After the supplementation, the dish was kept at room temperature for another day.

### Mass Spectrometry

Liquid NGM was prepared like solid NGM but without agar to prevent solidification. NGM was poured into petri dishes (7 ml NGM/ 6 cm petri dish) and seeded with bacteria (200 µl, OD_595_ = 0.9) or LB-medium (control). The medium was incubated for two days at room temperature to allow the bacteria to grow. Then, the bacteria were removed by centrifugation (4500 rpm, 10 min) and the medium was filtered (22 µm filter, Carl Roth P666.1) before using it for mass spectrometry analysis. Electrospray ionization mass spectrometry (ESI-MS) was used. For the LC-MS measurements, the liquid NGM samples were diluted 1:20 with methanol before the injection of 10 μl sample volumes. Triterpenes were separated on a Dionex HPG 3200 HPLC system (Thermo Scientific) equipped with a 150 × 2.1 mm, 2.7 μm, C18-CSH column (Waters) with a binary gradient system. Mobile phase A consisted of water + 0.1 % formic acid (FA), and mobile phase B consisted of methanol + 0.1 % FA. The mobile phase gradient was as follows: Starting conditions were 5 % mobile phase B, increased to 95 % B within 10 min, the plateau was held for 4 min, and the system was returned to starting conditions within 1 min and held for another 4.5 min. The flow rate was 0.5 mL/min. The MS and MS/MS analysis were performed with a quadrupole-time-of-flight instrument (maXis 4G, Bruker Daltonics, Bremen, Germany) equipped with an ESI source. The device was operated in positive-ion and negative-ion mode and the operating conditions were as follows: dry gas (nitrogen): 8.0 L/min, dry heater: 220 °C, nebulizer pressure: 1.8 bar, capillary voltage: 4500 V. Data analysis were performed using the software data analysis 4.2 and Metabolite detect 2.1 (Bruker daltonics, Bremen, Germany).

### Statistical analysis

If not stated, statistical analysis was performed in GraphPad Prism 6. For life-/healthspan assays, statistical analysis was done using OASIS (65, 66).

## Acknowledgments

Most nematode strains utilized in this work were provided by the Caenorhabditis Genetics Center, funded by the NIH Office of Research Infrastructure Programs (P40 OD010440). Other nematodes or bacteria strains were kindly provided by David Depomerai (University of Nottingham), Sheila Nathan (University of Malaysia), and Shawn Xu (University of Michigan). We thank Anne Hemmers, Bo Scherer, and Lisa Tschage for technical help, Sabine Metzger (University of Cologne) for the performance of mass spectrometry. We further thank Wormbase and the GENiE network funded by the European Cooperation in Science and Technology (COST Action BM1408). This work was financially supported by funding to N.V. from the German Research Foundation (DFG VE366/8-1). V.B. was supported by a Ph.D. scholarship by the Jürgen Manchot Foundation.

## Supplementary Information Text Methods

### Transformation of *E. coli* OP50(xu363)

For this study, we created OP50(L4440) by PEG transformation of OP50(xu363) according to methods described previously with adjustments (79). 100 µl of OP50(xu363) were transformed with 100 pg of plasmids isolated from the HT115(DE3) strains (Ahringer Library). The QIAprep Spin Miniprep Kit (Qiagen, 27104) was used for plasmid isolation.

### Crossing CZ2485 with transgenic strains

CZ2485 males were generated by treating hermaphrodite L4 larvae with heat stress (4 h at 30 °C). 10 adult CZ2485 males were then left with one hermaphrodite L3/L4 larvae for mating (P0 generation). Single worms of the F1 and F2 generations of this cross were checked for the *ju145* allele by PCR (ahr-1F CGGAAAGTTGATGTCTCTAC, ahr-1R TGCTGACTAGACGATATACC) followed by a restriction with *AlwI* (New England Biolabs, R0513S), which cuts the PCR fragment at the position of the *ju145* point mutation. Single worm PCR was performed according to standard protocols (1).

### Quantification of transgene expression

The expression of fluorescently-tagged genes was investigated in adult worms (first day of adulthood) by using fluorescence microscopy (ZEISS Imager M2 with an Axiocam MRm camera). For this, worms expressing fluorescently tagged genes were paralyzed with 15 mM NaN_3_ and pictures were taken with identical exposure times and settings. The pictures were then analyzed in either Image J, Fiji (2) or cellprofiler. Depending on the distribution of the expression, the fluorescence was either measured in a defined area of the region of interest or the whole worm. Statistical analysis was performed with the pooled data.

### Measurement of *cyp-35B1p::GFP* intensity in response to BaP

3-days old worms were treated with BaP or UVB for 18 hours. The relative intensity of *cyp-35B1p::GFP* was then visualized by fluorescence microscopy (100x magnification) and analyzed using Fiji (2). The integrated density was chosen as a parameter for the expression. The quantified intensities were normalized to the mean of untreated wild-type in each experiment. Statistical analysis was performed with the pooled data.

### UVB-induced Apoptosis

To investigate UVB-induced apoptosis, L4 larvae were treated with 600 J/m^2^ UVB and the apoptotic corpses were counted 24 h post-irradiation in the gonad loop region. The apoptotic corpses were identified based on their shape.

### Fertility

The number of eggs and progeny of animals were investigated during the main fertile period. Two days after synchronization single L4 larvae were transferred to NGM plates and from then transferred to fresh NGM plates every 24 hours until the 8^th^ day after hatching. The number of eggs laid during this period was counted. Two days later the progeny hatched on each day were counted.

### Pharyngeal pumping using the NemaMetrix ScreenChip System

To measure the pharyngeal pumping rate with the NemaMetrix ScreenChip, worms were washed off the plates with S-basal and collected in a reaction tube. Then, the worms were washed twice with S-basal and twice with 10 mM serotonin (Sigma Aldrich, 14927) and incubated in 10 mM serotonin for 30 min. Worms were loaded on the ScreenChip SC40 with a syringe (0.01 ml – 1 ml). The EPG of single worms was recorded for a duration of approx. 2 minutes. Only worms which showed pumping activity were recorded, while those with no pumping activity were not considered. The following NemAcquire-2.1 and NemAnalysis-0.2 software were used for analysis (https://nemametrix.com/products/software/).

### Assessment of mRNA expression by RTqPCR

Samples from 3 independent replicates with approximately 1000 3-days old worms per condition were collected and RNA was extracted. After washing and elution steps the RNA content was quantified by spectrophotometry, and 1 - 2 µg of RNA was used for the cDNA synthesis (Omniscript RT Kit (Qiagen, 205111). Primers were designed using NCBI Primer BLAST (https://www.ncbi.nlm.nih.gov/tools/primer-blast/) (3). Each primer pair was designed to span an exon-exon junction. Primer pairs and their features are listed in **Table S5**. For the Real-time qPCR, the cDNA was diluted 1:20 in 10 mM TRIS (pH 8.0). For the reaction, the GoTaq® qPCR kit (Promega, A6001) was used. The samples were run in a MyiQ2 cycler (BioRad), and the expression of each sample was measured in duplicate on the same multi-well plate. The expression was calculated relative to the reference genes *act-1* and *cdc-42* using the iQ5 software. All data collected were enabled for gene study according to the BioRad user instructions, and the expression was calculated using the normalized expression (ddC_T_). The efficiency of each primer pair reaction was added for correct quantification of the normalized expression. The efficiency was assessed with 1:20, 1:100, 1:500, and 1:2500 dilutions of the cDNA. From normalized expression values, the fold-change compared to wild-type was calculated for each replicate.

---

**Fig. S1.**
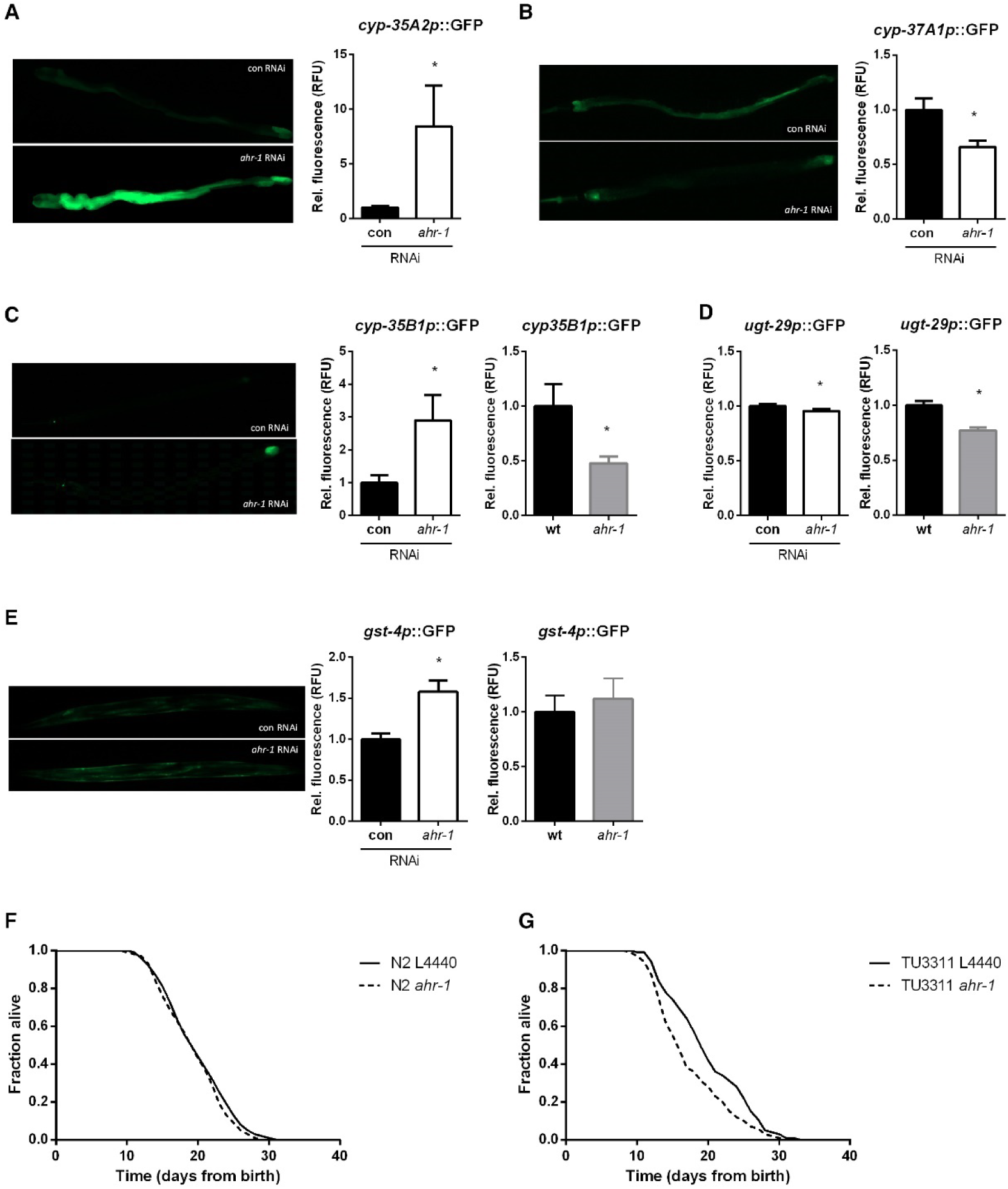
AHR-1 regulates genes involved in xenobiotic stress response. A) Quantification of cyp-35A2p::GFP expression in worms treated with control or ahr-1 RNAi. Mean + 95 % CI of pooled data from 3 independent experiments with 43 worms are shown. B) Quantification of cyp-37A1p::GFP expression in worms treated with control or ahr-1 RNAi. Mean + 95 % CI of pooled data from 2 independent experiments with 38 (con RNAi) and 26 (ahr-1 RNAi) worms are shown. C) Quantification of cyp-35B1p::GFP expression in worms treated with control or ahr-1 RNAi (left panel) or worms with either wildtype ahr-1 or the ju145 allele (right panel). Mean + 95 % CI of pooled data from 60 (con RNAi), 40 (ahr-1 RNAi), 52 (wt), and 63 (ahr-1) worms in 3 independent experiments are shown. D) Quantification of ugt 29p::GFP expression in ahr-1-depleted worms. Mean + 95 % CI of pooled data from 3 independent experiments are shown. Number of worms 44 (con RNAi), 36 (ahr-1 RNAi) [left panel], 34 (wt), 40 (ahr-1) [right panel]. E) Quantification of gst 4p::GFP expression in worms treated with control or ahr-1 RNAi (left panel) or worms with either wildtype ahr-1 or the ju145 allele (right panel). Mean + 95 % CI of pooled data from 27 (con RNAi), 29 (ahr-1 RNAi), 27 (wt), and 19 (ahr-1) worms in 2 independent experiments are shown. F) ahr-1 depletion via RNAi does not affect the lifespan of N2. Kaplan Meier curves of control (L4440) or ahr-1 RNAi treated worms. Pooled data of 280 (N2 L4440) and 264 (N2 ahr-1) worms/condition in 5 independent replicates are shown. * p-value < 0.05 vs. wt, statistical test: Log-rank test. G) ahr-1 depletion via RNAi shortens the lifespan in a strain with enhanced RNAi in the nervous system (TU3311). Kaplan Meier curves of control (L4440) or ahr-1 RNAi treated worms. Pooled data of 180 worms/condition in 3 independent replicates are shown. * p-value < 0.05 vs. wt, statistical test: Log-rank test.

**Fig. S2.**
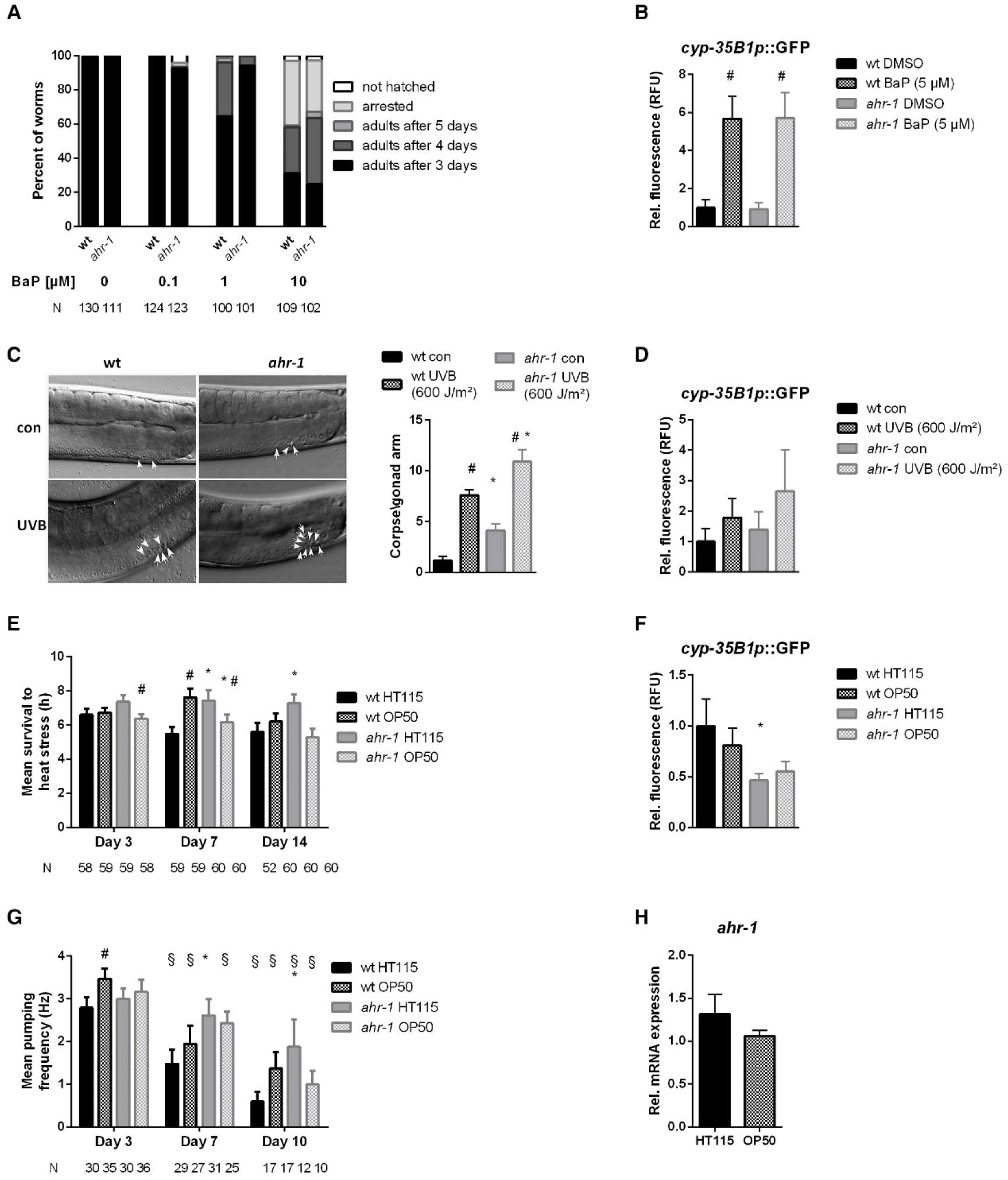
Aging is influenced by the bacterial diet in the *ahr-1* mutant. A) Development of wild-type and *ahr-1* on indicated doses of BaP. Pooled data of 3 independent experiments are shown, the number of individuals is presented as N. Statistical test: 2-Way ANOVA with Tukey’s multiple comparisons test, no statistical differences were observed. B) cyp-35B1p::GFP induction in response to 5 µM of BaP. Mean + 95 % CI of pooled data of 58 (wt DMSO), 51 (wt BaP), 53 (ahr-1 DMSO), and 65 (ahr-1 BaP) in 3 independent experiments are shown. * p-value < 0.05 vs. wt, # p-value < 0.05 vs. DMSO, statistical test: One-way ANOVA with Tukey’s multiple comparisons test. C) UVB irradiation and ahr-1 loss of function induce apoptosis. Mean + 95 % CI of pooled data of 39 (wt con, ahr-1 UVB) and 38 (wt UVB, ahr-1 con) worms in 3 independent experiments are shown. Statistical test: 2-Way ANOVA with Tukey’s multiple comparisons test, * p-value < 0.05 vs. wt, # p-value < 0.05 vs. con. D) cyp-35B1p::GFP induction in response to UVB irradiation. Mean + 95 % CI of pooled data of 34 (wt con), 37 (wt UVB), 28 (ahr-1 con), and 35 (ahr-1 UVB) worms in 2 independent experiments are shown. * p-value < 0.05 vs. wt, # p-value < 0.05 vs. control, statistical test: One-way ANOVA with Tukey’s multiple comparisons test. E) Heat shock resistance of wild-type and ahr-1 feeding on HT115 or OP50. Mean survival times + SEM of 3 independent experiments are shown, the number of individuals is presented as N. Statistical test: Two-way ANOVA with Tukey’s multiple comparisons test. * p-value < 0.05 vs. wt, # p-value < 0.05 vs. HT115, F) cyp-35B1p::GFP expression of worms feeding on HT115 or OP50. Mean + 95 % CI of pooled data of 37 (wt HT115), 27 (wt OP50), 44 (ahr-1 HT115), and 19 (ahr-1 OP50) in 2 independent experiments are shown. * p-value < 0.05 vs. wt, # p-value < 0.05 vs. HT115, statistical test: One-way ANOVA with Tukey’s multiple comparisons test. G) Pumping frequencies of pooled data from 3 independent experiments are shown as mean + 95 % CI, the number of individuals is presented as N. * p-value < 0.05 vs. wt, # p-value < 0.05 vs. HT115, § p-value < 0.05 vs. day 3, statistical test: Two-way ANOVA with Tukey’s multiple comparisons test. H) ahr-1 mRNA expression in wild-type worms feeding on HT115 or OP50. Pooled data of 3 independent replicates are shown. No statistical significance was observed with the t-test.

**Fig. S3.**
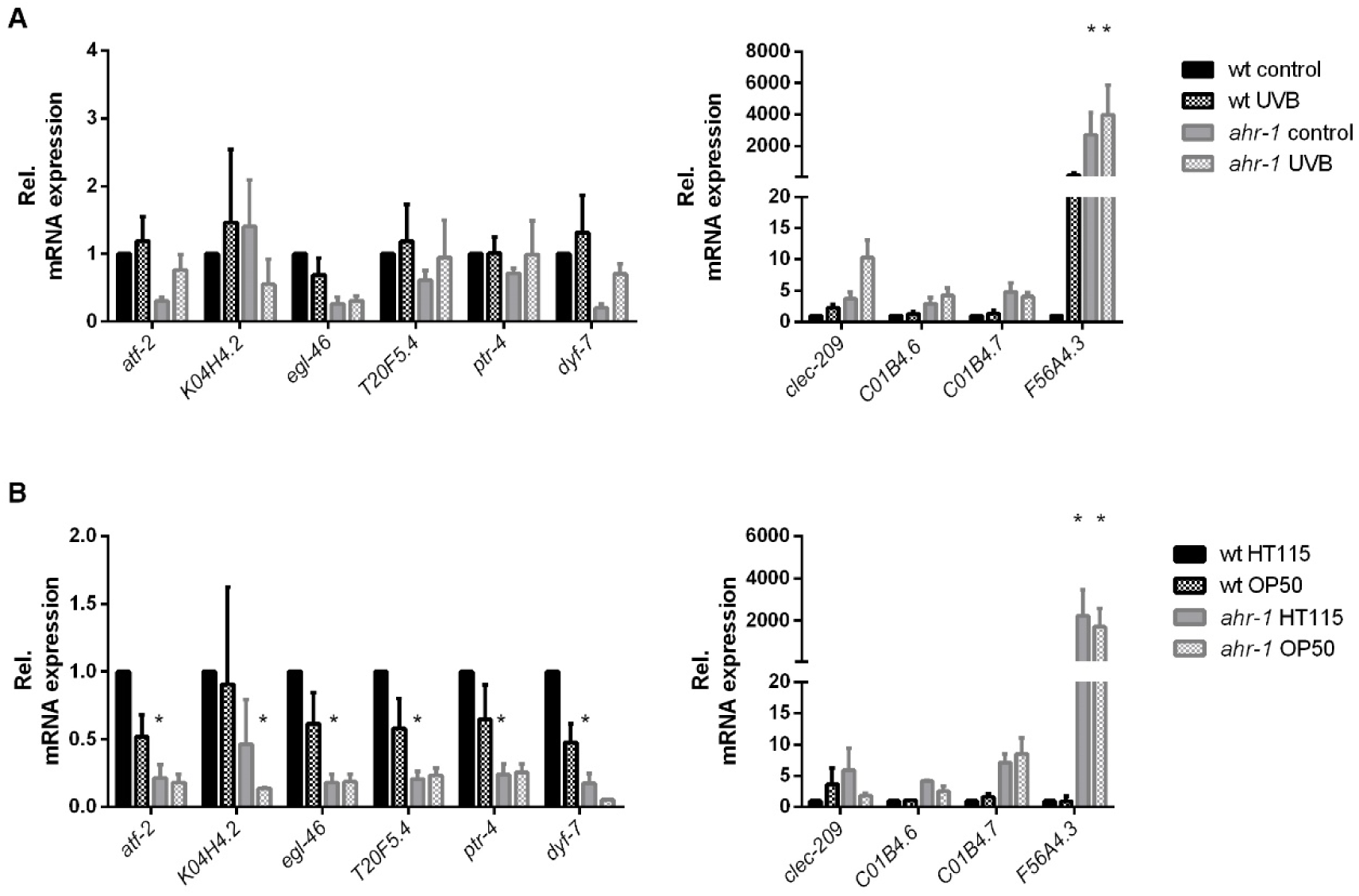
Gene expression changes in response to Ahr modulators. Expression of the strongest up- and down-regulated genes in *ahr-1* vs. wild-type in response to A) 1200 J/m^2^ UVB, and B) bacteria. Pooled data from 3 independent replicates are shown. Statistical test: 2-Way ANOVA with Tukey’s multiple comparisons test, * significance vs. wt, # significance vs. control.

**Fig. S4.**
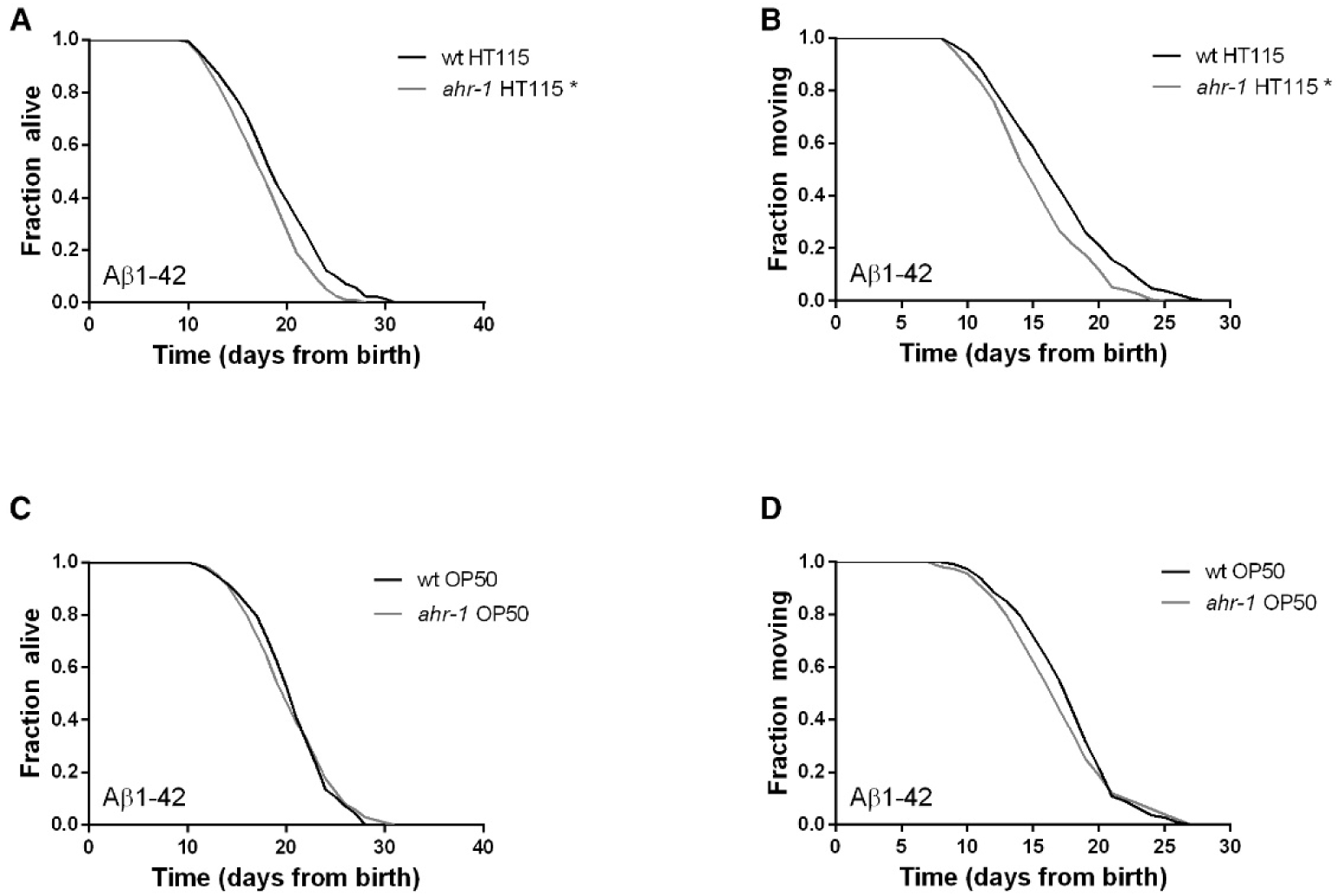
AHR-1 does not influence aging in an Alzheimer’s disease model. A – B) Kaplan Meier curves of Abeta;wt and Abeta;ahr-1 grown on HT115. Pooled data of 240 (wt) and 236 (*ahr-1*) worms/condition in 4 independent replicates are shown. * p-value < 0.05 vs. wt, statistical test: Log-rank test. C – D) Kaplan Meier curves of Abeta;wt and Abeta;ahr-1 grown on OP50. Pooled data of 120 worms/condition in 2 independent replicates are shown. Statistical evaluation with log-rank test did not display significant differences.

**Fig. S5.**
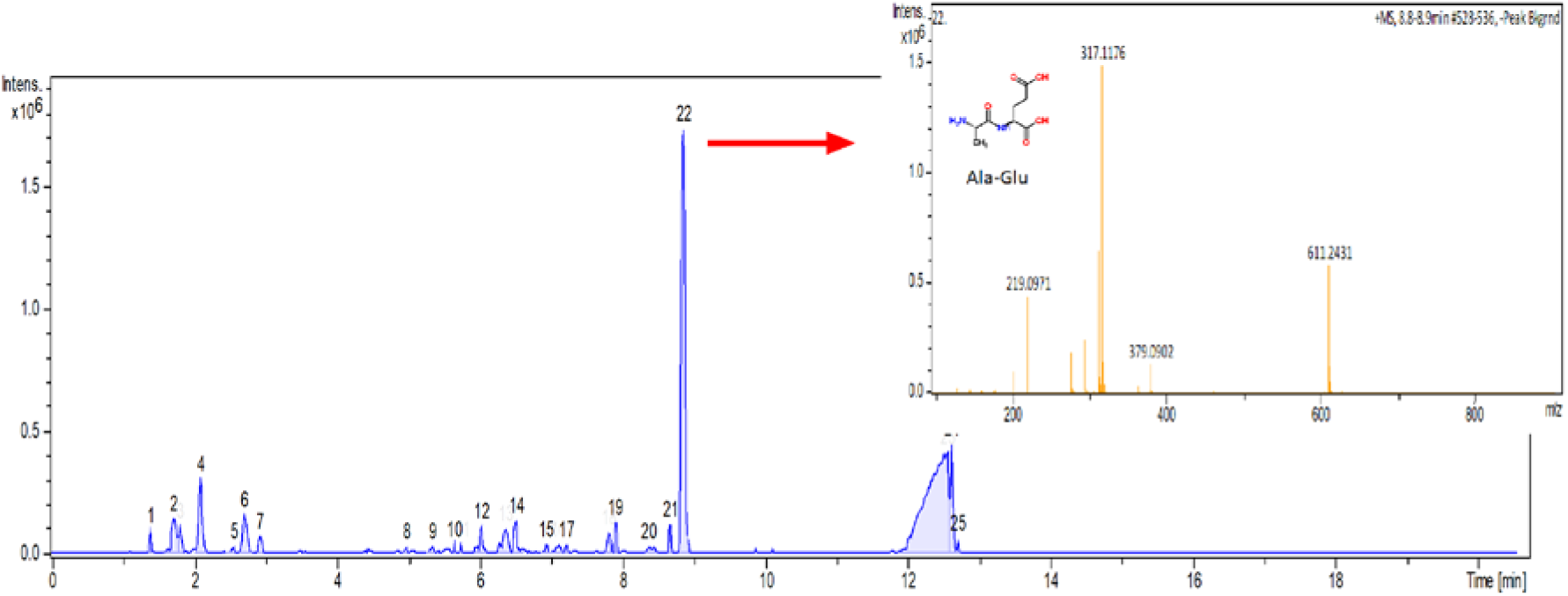
Positive ESI-mass spectrum of OP50(L4440) medium after subtraction of the HT115(L4440) spectrum. Masses peak 22 are shown as inset. m/z 219 is likely an Ala-Glu dipeptide.

**Table S1.** Relative expression of transgenes in control or *ahr-1* RNAi-treated reporter strains. The relative expression was measured in worms on their first day of adulthood and normalized to control RNAi treated worms in each replicate. N shows the number of worms and n the number of experiments, Statistical test: 2-tailed unpaired t-test.

| **Strain name** | **Gene** | **Rel. expression**<br>**(mean ± SD)** | **N** | **n** | **p-value** |
| --- | --- | --- | --- | --- | --- |
| BC14926 | ***cyp-14A3*** | anterior: 0.97 | 38 (con) | 3 | 0.544 |
|  |  | posterior: 1.05 | 35 (*ahr-1*) |  | 0.333 |
| SD1444 | ***cyp-25A2*** | 1.07 ± 0.16 | 38 | 2 | 0.067 |
| BC20334 | ***cyp-29A2*** | 0.85 ± 0.60 | 42 | 2 | 0.377 |
| cyp-35A2 | ***cyp-35A2*** | anterior: 8.43 ± 11.96 | 43 (con) | 3 | < 0.001 |
| | | posterior: 1.99 ± 2.14 | 42 (*ahr-1*) | | 0.005 |
| cyp-35A3 | ***cyp-35A3*** | 1.08 ± 0.09 | 28 (con)<br>26 (*ahr-1*) | 2 | 0.02 |
| CY573 | ***cyp-35B1*** | anterior: 2.60 ± 1.97 | 36 (con)<br>27 (*ahr-1*) | 2 | < 0.001 |
| | | posterior: 2.90 ± 2.40 | 55 (con)<br>44 (*ahr-1*) | 3 | < 0.001 |
| BC15044 | ***cyp-37A1*** | 0.66 ± 0.14 | 46 (con)<br>26 (*ahr-1*) | 2 | < 0.001 |
| CL2166 | ***gst-4*** | 1.58 ± 0.37 | 27 (con)<br>29 (*ahr-1*) | 2 | < 0.001 |
| ugt-29 | ***ugt-29*** | 0.96 ± 0.05 | 44 (con)<br>36 (*ahr-1*) | 3 | 0.004 |

**Table S2.**
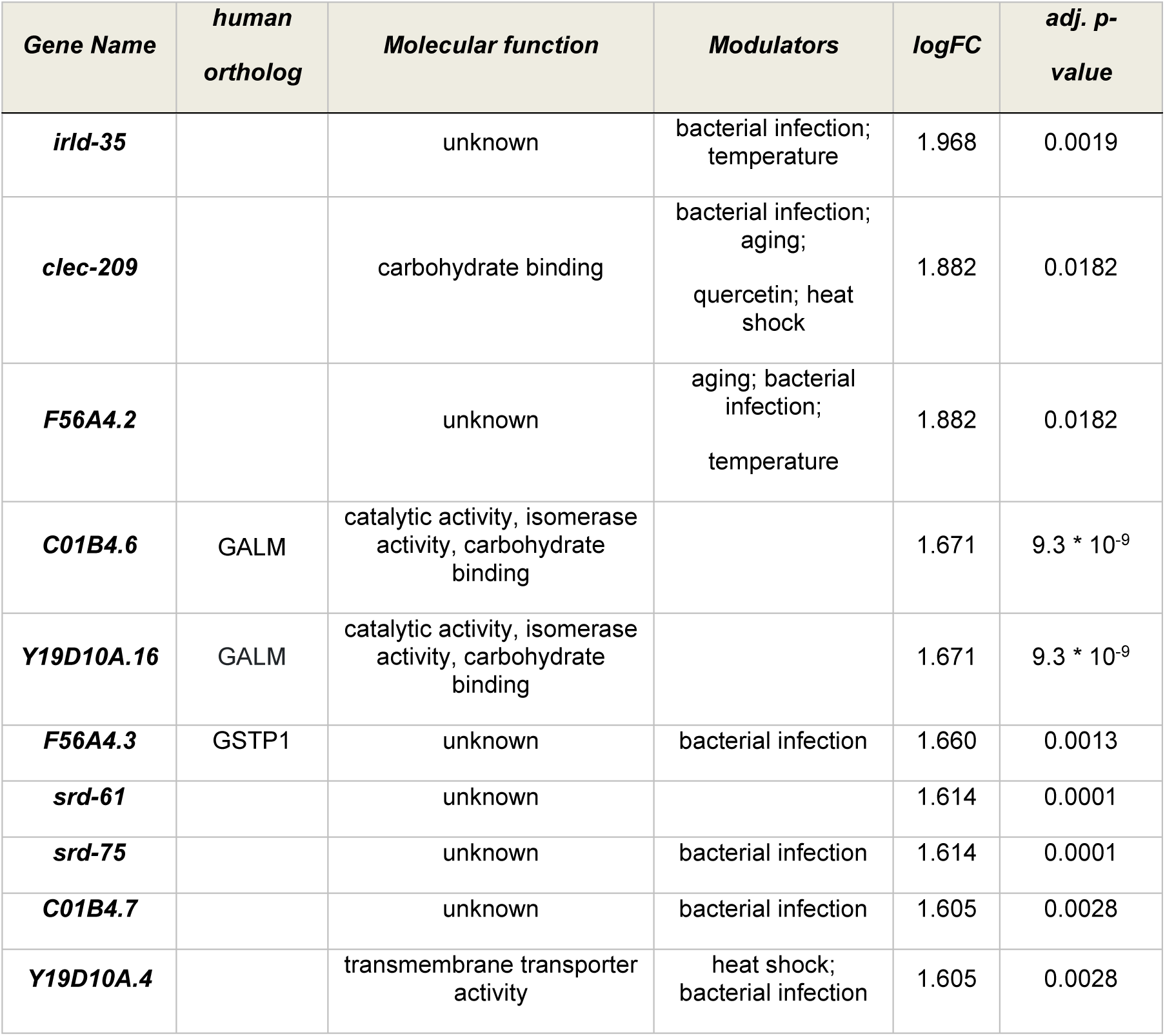
List of the most robust over-expressed genes in *ahr-1 vs*. wildtype. Human orthologs were extracted using the BioMart tool (https://parasite.wormbase.org/biomart). Modulators of these genes with potential relevance for AHR-1 activity were selected from WormBase. For log fold change values (logFC) a base of 2 is used.

**Table S3.**
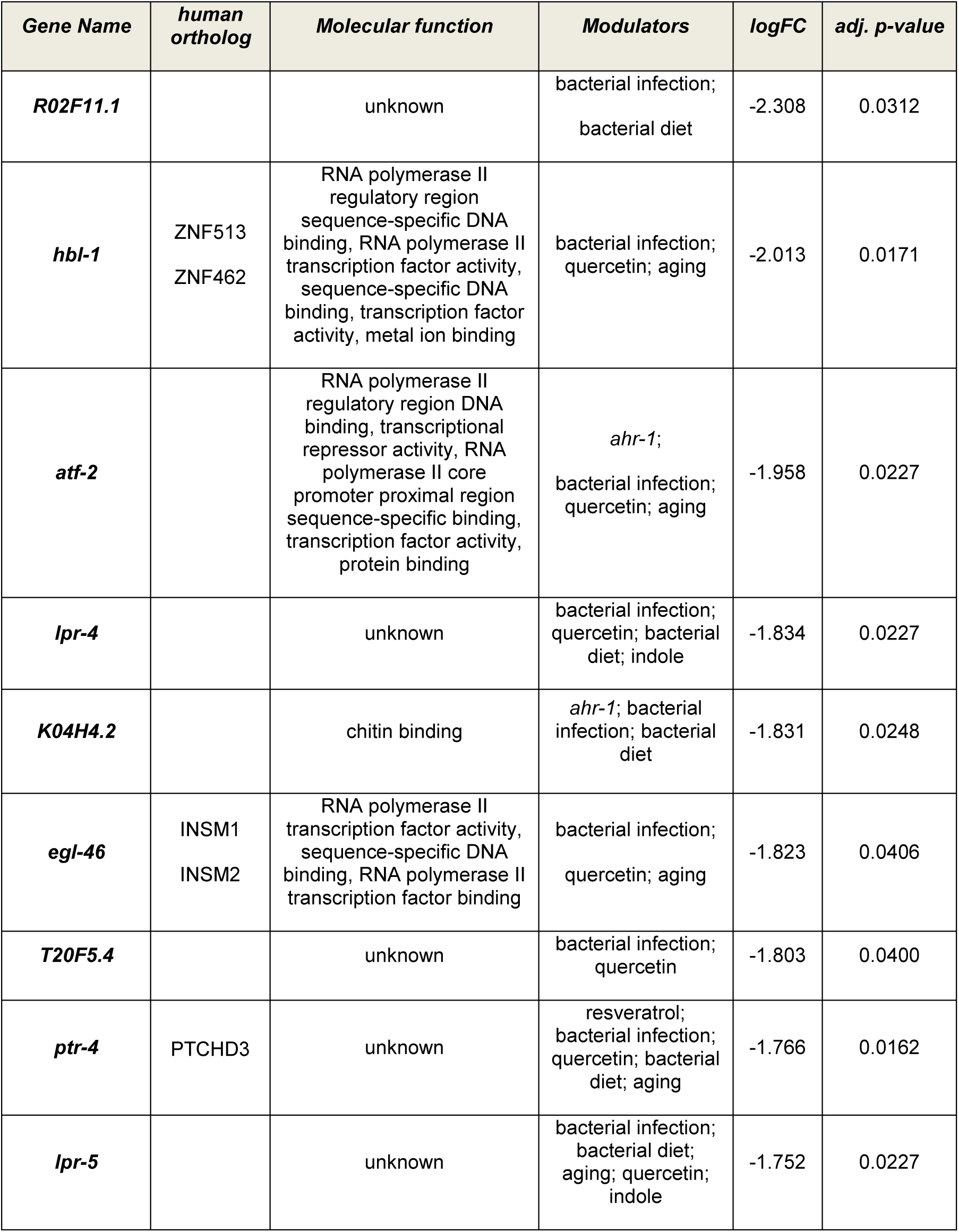
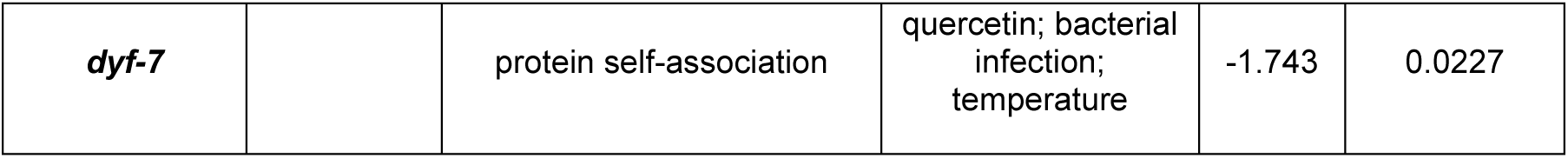
List of the most robust under-expressed genes in *ahr-1* vs. wildtype. Human orthologs were extracted using the BioMart tool (https://parasite.wormbase.org/biomart). Modulators of these genes with potential relevance for AHR-1 activity were selected from WormBase. For log fold change values (logFC) a base of 2 is used.

| **Gene Name** | **human ortholog** | **Molecular function** | **Modulators** | **logFC** | **adj. p-value** |
| --- | --- | --- | --- | --- | --- |
| ***R02F11.1*** |  | unknown | bacterial infection;<br>bacterial diet | -2.308 | 0.0312 |
| ***hbl-1*** | ZNF513<br>ZNF462 | RNA polymerase II regulatory region sequence-specific DNA binding, RNA polymerase II transcription factor activity, sequence-specific DNA binding, transcription factor activity, metal ion binding | bacterial infection;<br>quercetin; aging | -2.013 | 0.0171 |
| ***atf-2*** |  | RNA polymerase II regulatory region DNA binding, transcriptional repressor activity, RNA polymerase II core promoter proximal region sequence-specific binding, transcription factor activity, protein binding | *ahr-1*;<br>bacterial infection;<br>quercetin; aging | -1.958 | 0.0227 |
| ***lpr-4*** |  | unknown | bacterial infection;<br>quercetin; bacterial diet; indole | -1.834 | 0.0227 |
| ***K04H4.2*** |  | chitin binding | *ahr-1*; bacterial infection; bacterial diet | -1.831 | 0.0248 |
| ***egl-46*** | INSM1<br>INSM2 | RNA polymerase II transcription factor activity, sequence-specific DNA binding, RNA polymerase II transcription factor binding | bacterial infection;<br>quercetin; aging | -1.823 | 0.0406 |
| ***T20F5.4*** |  | unknown | bacterial infection;<br>quercetin | -1.803 | 0.0400 |
| ***ptr-4*** | PTCHD3 | unknown | resveratrol;<br>bacterial infection;<br>quercetin; bacterial diet; aging | -1.766 | 0.0162 |
| ***lpr-5*** |  | unknown | bacterial infection;<br>bacterial diet;<br>aging; quercetin;<br>indole | -1.752 | 0.0227 |
| ***dyf-7*** |  | protein self-association | quercetin; bacterial infection; temperature | -1.743 | 0.0227 |

**Table S4.**
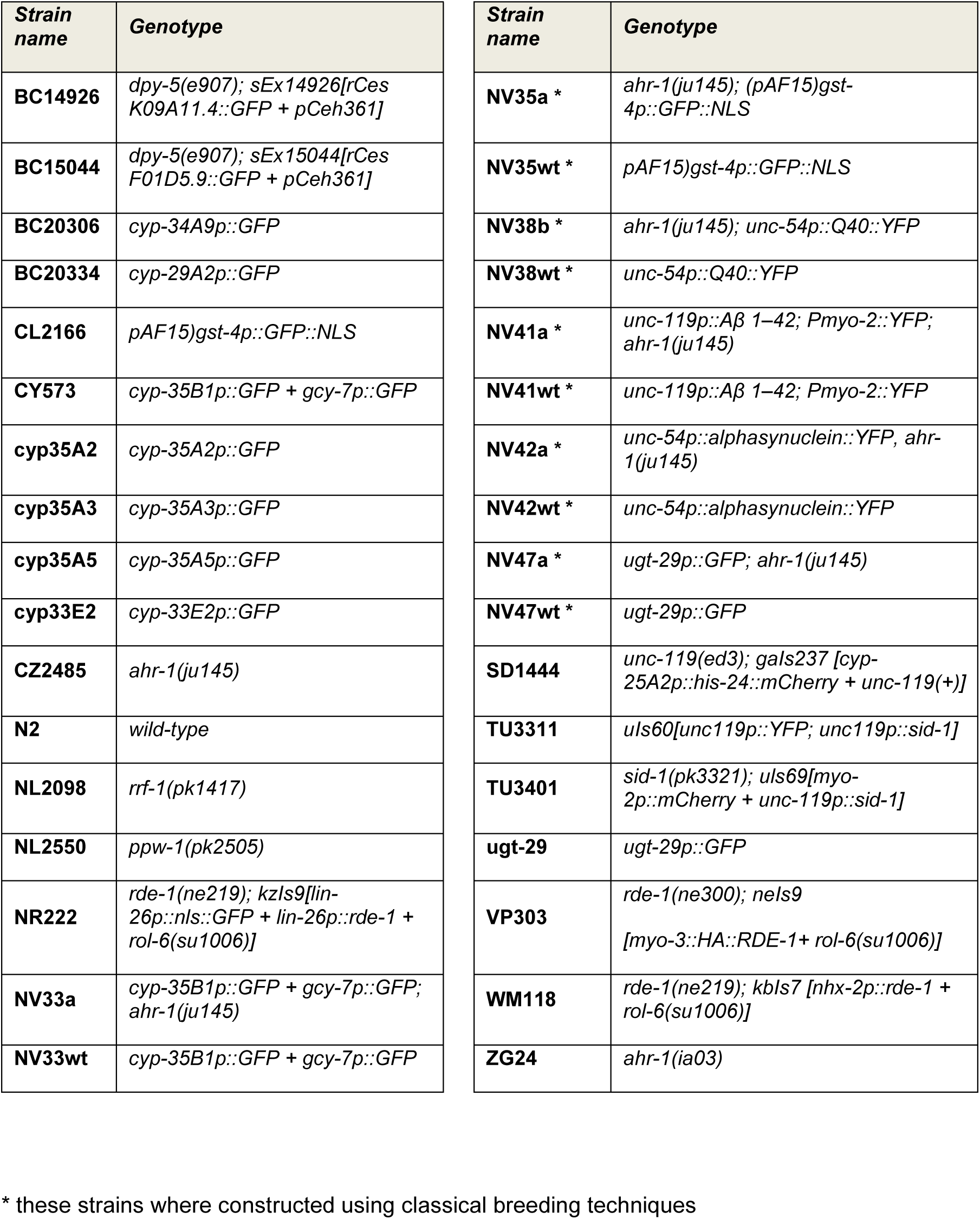
*C. elegans* strains used in this study. Strains are sorted alphabetically.

**Table S5.**
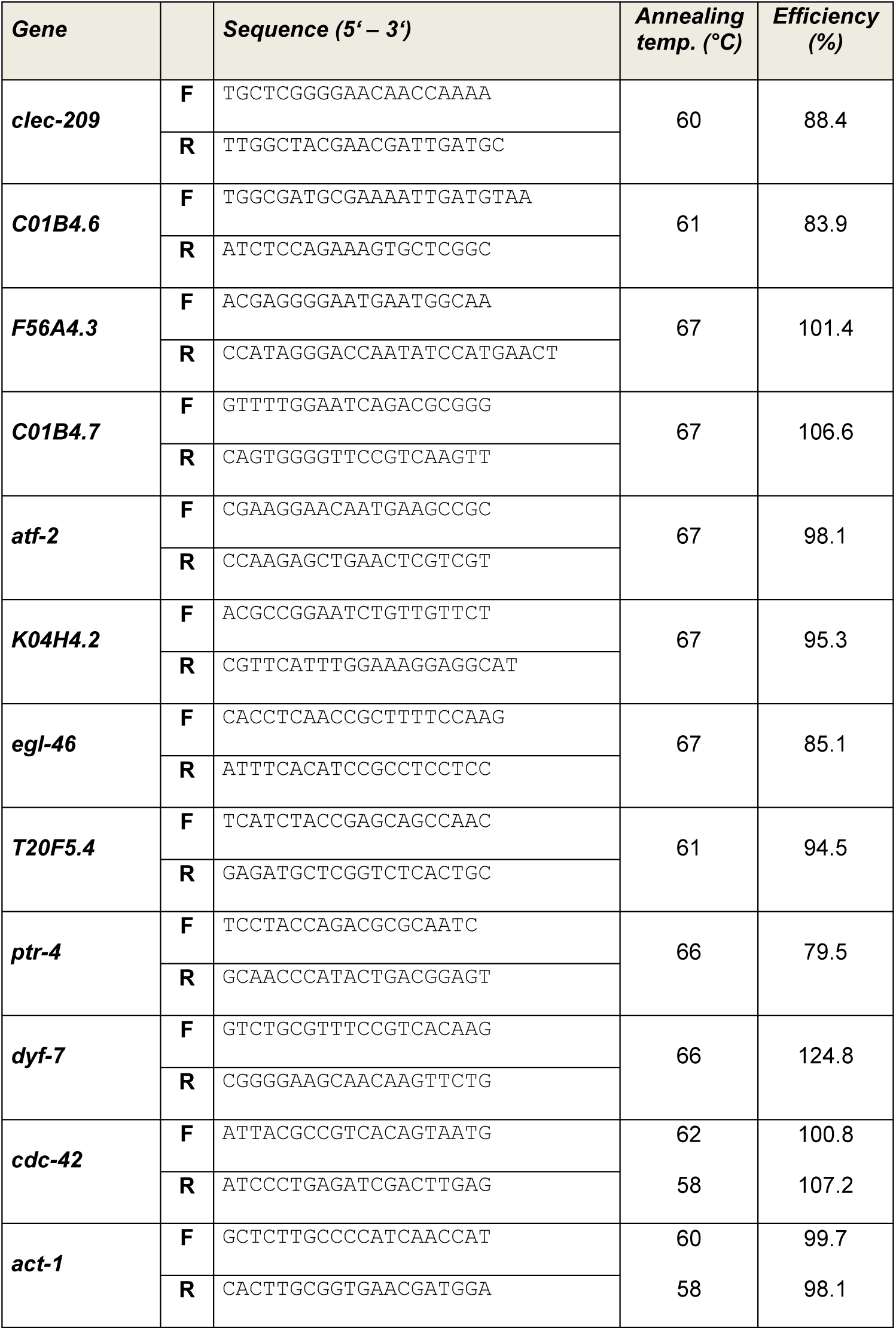
Primer pairs used in this study to validate genes differentially expressed between *ahr-1(ju145)* and wild-type (Fig. S3).

